# Enhancing the Fitness of Embryoid Bodies and Organoids by Chemical Cytoprotection

**DOI:** 10.1101/2022.03.21.485225

**Authors:** Seungmi Ryu, Claire Weber, Pei-Hsuan Chu, Carlos A. Tristan, Ben Ernest, Vukasin M. Jovanovic, Tao Deng, Jaroslav Slamecka, Hyenjong Hong, John Braisted, Marissa Hirst, Anton Simeonov, Ty C. Voss, Ilyas Singeç

## Abstract

Embryoid bodies (EBs) and self-organizing organoids derived from human pluripotent stem cells (hPSCs) recapitulate tissue development in a dish and hold great promise for disease modeling and drug development. However, current protocols are hampered by cellular stress and apoptosis during cell aggregation, resulting in variability and impaired cell differentiation. Here, we demonstrate that EBs and various organoid models (e.g., brain, gut, and kidney) can be optimized by using the CEPT small molecule cocktail, a polypharmacology approach that ensures cytoprotection and cell survival. Application of CEPT (chroman 1, emricasan, polyamines, trans-ISRIB) for just 24 hours during cell aggregation has long-lasting consequences affecting morphogenesis, gene expression, and cellular differentiation. Various qualification methods confirmed that CEPT treatment consistently improved EB and organoid fitness as compared to the widely used ROCK inhibitor Y-27632. Collectively, we discovered that stress-free cell aggregation and superior cell survival in the presence of CEPT are critical quality control determinants that establish a robust foundation for bioengineering complex tissue and organ models.

## INTRODUCTION

Controlling cell fate, differentiation, and maturation of human tissues *in vitro* are among the most formidable challenges in biomedical research. Self-renewing hPSCs, such as induced pluripotent stem cells (iPSCs), serve as an inexhaustible source of human tissues that provide valuable insights into normal development and human diseases (Eiraku et al., 2008; Lancaster and Huch, 2019; Mansour et al., 2018; Sato et al., 2009; Velasco et al., 2019). *In vitro*-generated organoids exhibit a remarkable potential for self-organization and recapitulate important aspects of organogenesis. Depending on the tissue of interest, different protocols have been developed to control lineage commitment and cell fate specification by modulating specific cell signaling pathways and providing the appropriate physicochemical environment for differentiating cells (Hendriks et al., 2020). Recent cell culture advances allow for extended culture of organoids for periods ranging from several months to years (Lee et al., 2020; Sloan et al., 2017; Velasco et al., 2019).

Stem cells are generally grown on cell culture plates coated with a substrate (e.g., laminin, vitronectin) and generation of three-dimensional (3D) cultures from these cells requires detachment and single-cell dissociation. Dissociated hPSCs respond with hyperactive cell contractions as one of the underlying mechanisms that lead to cell death, which is typically blocked by applying 10-50 µM Y-27632 (Ohgushi et al., 2010; Watanabe et al., 2007, Koehler et al., 2017; Lancaster et al., 2018; Mansour et al., 2018; Velasco et al., 2019). Preparing the cellular material, plating cells into ultra-low attachment plates, and aggregating cells into 3D structures are associated with marked cellular stress and cell death by apoptosis and anoikis (Chen et al., 2021). To deal with poor cell survival, one can compensate by plating a higher number of cells or by using reagents that partially improve cell survival such as Y-27632. Indeed, suboptimal approaches can contribute to product variation and non-standardized organoid models that pose challenges for disease modeling and drug discovery. Variables that affect the quality of 3D cultures include the cell numbers plated, cell survival during initial cell aggregation and long-term culture, variable differentiation propensity of different cell lines, and other factors that contribute to biological and technical variability (Ortmann and Vallier, 2017). In the present study, we performed various systematic analyses and provide evidence that the CEPT cocktail improves the formation, proper differentiation, and overall reproducibility of EBs and organoids of different developmental lineages.

## RESULTS

### Optimizing EB formation using CEPT

EB formation is a widely used assay to measure the pluripotent differentiation potential of hPSCs (Keller, 1995; Tsankov et al., 2015). Differentiating EBs emulate the gastrulation process of the developing embryo and generate the three primary germ layers (ectoderm, mesoderm, endoderm). EB formation is often also the first step when generating various organoid models from hPSCs (Eiraku et al., 2008; Gabriel et al., 2021; Lancaster and Knoblich, 2014; Quadrato et al., 2017; Saha et al., 2022). For instance, a kit-based brain organoid protocol (STEMCELL Technologies) uses EB formation as the first step (**Figure 1A**). We first confirmed that the recently developed CEPT cocktail was superior to Y-27632 or when only two components (chroman 1 + emricasan; “C+E”) of the cocktail were used. When varying cell densities, the use of CEPT consistently yielded higher cell numbers (CellTiter-Glo assay) and larger EBs as measured at day 1 and day 5 (**Figures 1B and 1C**). When plating the same cell numbers, the diameter of EBs was consistently larger after CEPT treatment versus Y-27632 (**Figure 1D**). Interestingly, EBs of comparable size were generated with CEPT and plating fewer cells (6,000 cells) versus Y-27632 requiring higher cell numbers (9,000 cells)(**Figure 1D**). Improved cell survival and EB formation due to CEPT was confirmed in hESCs (WA09) and three different iPSC lines (LiPSC-GR1.1, GM23279, GM25256)(**Figures S1A and S1B**). Moreover, improved EB formation by CEPT was confirmed using live-cell microscopy (day 1 and 5) and the application of dyes that label live cells (calcein-AM) and dead cells (EthD-1)(**Figure 1E**). Confocal microscopy underscored the importance of using CEPT for optimal cell aggregation during the first 24 hours (**Figure S1C**). In contrast, Y-27632 treatment resulted in dead cells that were not only attached to the EB surface but also found to be enclosed within the EB as revealed by reconstructing confocal sections (Z-stack). This suggests that dead cells and debris become part of EBs during cell aggregation when suboptimal conditions are employed.

**Figure 1.**
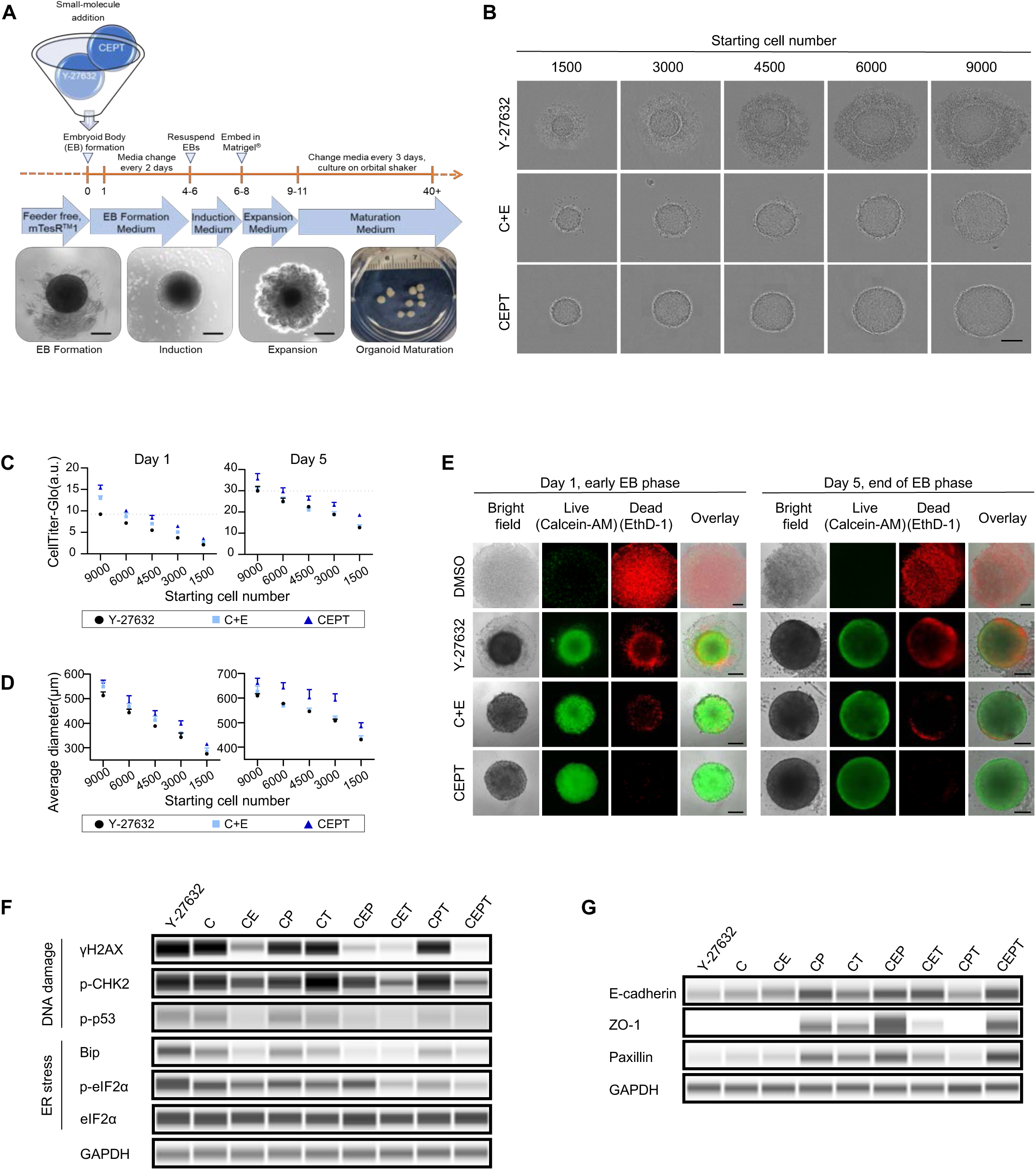
Improved cell viability and EB morphology by CEPT. (A) Overview of the protocol that was used to compare the effects of Y-27632 and the CEPT cocktail on EBs and brain organoids. (B) Morphology of single EBs in 96-well ultra-low attachment plates after plating different cell numbers and treatment with Y-27632 and CEPT. Images were taken 24 hours after plating cells. (C, D) Quantitative analysis of single EBs measuring viability and size. Cell viability was measured using the CellTiter-Glo assay (ATP levels) and EB diameter was determined using high-content imaging (Celigo). (E) Visualization of live and dead cells in single EBs after treatment with Y-27632 and CEPT for 24 hours. Cells were stained with calcein-AM for live (green) and EthD-1 for dead cells (red). Images were taken on day 1 and 5. (F) Western blot analysis showing cellular stress markers upon EB formation at 3 hours after small molecule compounds addition. GAPDH was used as a loading control. (G) Western blot analysis showing cell adhesion molecules expressed in the formed EB. GAPDH was used as a loading control. All scale bars, 200 µm.

Next, we asked how Y-27632 and different combinations of the components of the CEPT cocktail may affect markers of DNA damage, endoplasmic reticulum (ER) stress, and the expression of cell membrane-associated proteins. Western blot experiments demonstrated that the application of CEPT resulted in the most favorable condition, which is avoidance of cellular stress, reduced DNA damage, and higher expression levels of E-cadherin, ZO-1, and paxillin (**Figures 1F and 1G**).

### CEPT enhances EB formation for high-throughput applications

To further evaluate the effects of improved cell survival and stress-free EB formation, we focused on analyzing single EBs. Profiling gene expression from single EBs is technically challenging since the RNA yield is typically very low and therefore difficult to implement for high-throughput experiments. To overcome this challenge, we established RNA-mediated oligonucleotide Annealing, Selection, and Ligation with next-generation sequencing (RASL-Seq) for analysis of single EBs in 384-well plates (**Figure 2A**). This targeted transcriptomic approach allows direct and reproducible analysis of small RNA amounts as little as 10 ng (Li et al., 2012). RASL-seq is an efficient method for large-scale transcriptomic analysis of single EBs since this technology enables direct analysis of RNA levels in EB lysates without the need for cDNA generation and can therefore be performed in a fully automated fashion (**Figure 2A**). To this end, we designed a probe set focusing on lineage-specific genes (19 genes for endoderm, 12 genes for mesoderm, 21 genes for ectoderm; genes listed in **Figure 2E**). Single EBs (20,000 cells/well) generated in the presence of Y-27632 or CEPT during the first 24 hours were cultured in E6 medium for 7 days and then processed for RASL-Seq analysis (**Figure 2A**). Gene expression profiling of single EBs (n = 16 per group) showed multi-lineage differentiation in both conditions (**Figures 2B and 2C**). However, when analyzing the biological variability across individual EBs by measuring the coefficient of determination (R^2^), which is a measure of the similarity across individual EBs, we observed considerable differences (**Figure 2D**). Indeed, the coefficient was higher after CEPT treatment versus Y-27632 (total of 240 paired values comparing 16 individual EBs within the same treatment group). While 17.5% of the paired comparisons among EBs generated with Y-27632 showed an R^2^ value lower than 0.8, the value for CEPT-treated EBs was only 9.17%. Variation in expression of each gene across individual EBs indicated as a coefficient of variation (CV), was higher in the Y-27632 treatment versus CEPT (**Figures 2E and 2F**) for important lineage-determining transcription factors (e.g., SOX17 for endoderm, HAND2 for mesoderm, and PAX6 for ectoderm). Collectively, these findings suggest that CEPT reduces the variability promotes standardization of EBs, an *in vitro* model that is widely used to study pluripotent differentiation potential and generate organoids of different developmental lineages.

**Figure 2.**
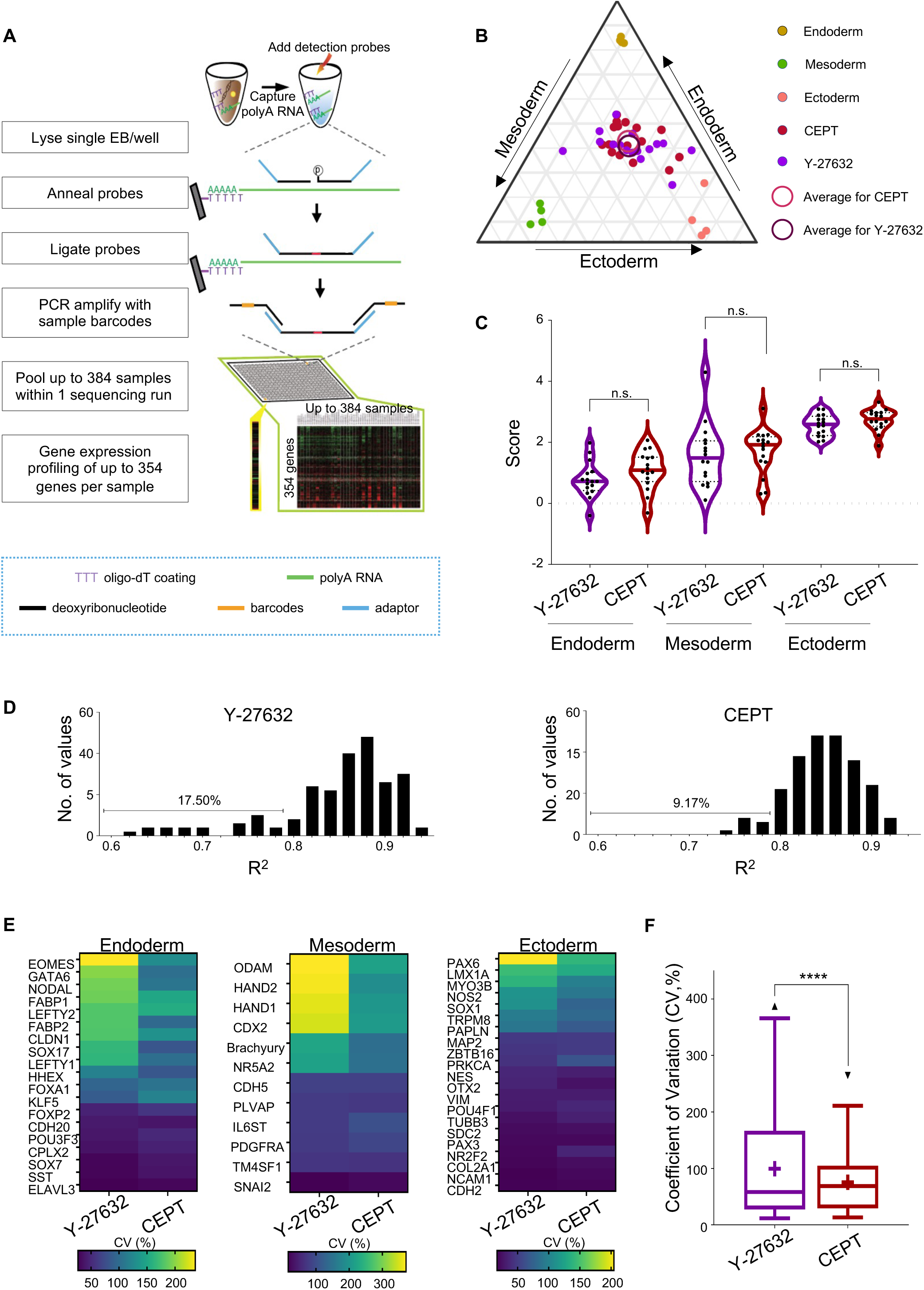
Effect of the CEPT cocktail on differentiation of individual EBs. (A) Overview of the RASL-Seq method enabling efficient targeted transcriptomics of single EBs. (B) Triangle plot comparing EB differentiation potential in chemically defined E6 medium after treatment with Y-27632 or CEPT. (C) Violin plot showing no significant difference between Y-27632 and CEPT-treated EBs regarding gene expression scores for endoderm, mesoderm, and ectoderm (see Supplemental Information for details). (D) Pearson correlation analysis and distribution plot of R^2^ values for individual EBs within each treatment group (Y-27632 versus CEPT). Note the overall distribution and occurrence of EBs with lower correlation (R^2^ < 0.8). The higher number in the Y-27632 group (17.5%) versus CEPT (9.17%) indicates relatively higher variability of EBs. (E) Heatmap analysis indicating more heterogeneous gene expression levels across individual EBs after Y-27632 treatment as shown by higher CV values as compared to CEPT. (F) Box and whisker plots of the compiled CV distribution. Boxes represent median, the whiskers denote minimum and maximum values, and + represents mean value. All data were obtained using RASL-seq analysis of single EBs (n = 16 per group) that were cultured in chemically defined E6 medium for 7 days.

### Optimizing brain organoids by using CEPT

To generate brain organoids, we used a kit-based method (**Figure 1A and 3A**) to form EBs, which then differentiate into neural structures with prominent neuroepithelial buds (Lancaster and Knoblich, 2014). These neuroepithelial buds represent neural tube-like structures and recapitulate some aspects of early brain development. We asked whether CEPT treatment might affect the morphogenesis of brain organoids. To this end, we compared organoids generated with Y-27632 or CEPT added for 24 hours during EB formation. By the end of the neuroepithelial expansion phase on day 10 (**Figure 3A**), whole organoids were fixed and subjected to optical clearing using a published protocol (Boutin et al., 2018). High-content confocal microscopy was used to generate images from individual organoids, which were stained with the nuclear marker Hoechst to visualize all cells (**Figure 3B**). Automated image analysis based on segmentation and 3D rendering was performed to detect and quantify the number of neural buds generated per 1,000 cells (**Figure 3C**), the density of buds per organoid volume (**Figure 3D**), and bud size (**Figure 3E**). This systematic analysis revealed that CEPT-generated organoids contained higher numbers of neuroepithelial buds as compared to Y-27632. We also noted that the size of the neuroepithelial buds correlated with the number of seeded cells in the CEPT treatment group (R^2^ = 0.8848), while such correlation was not found for organoids generated using Y-27632 (R^2^ = 0.4178)(**Figure 3E**).

**Figure 3.**
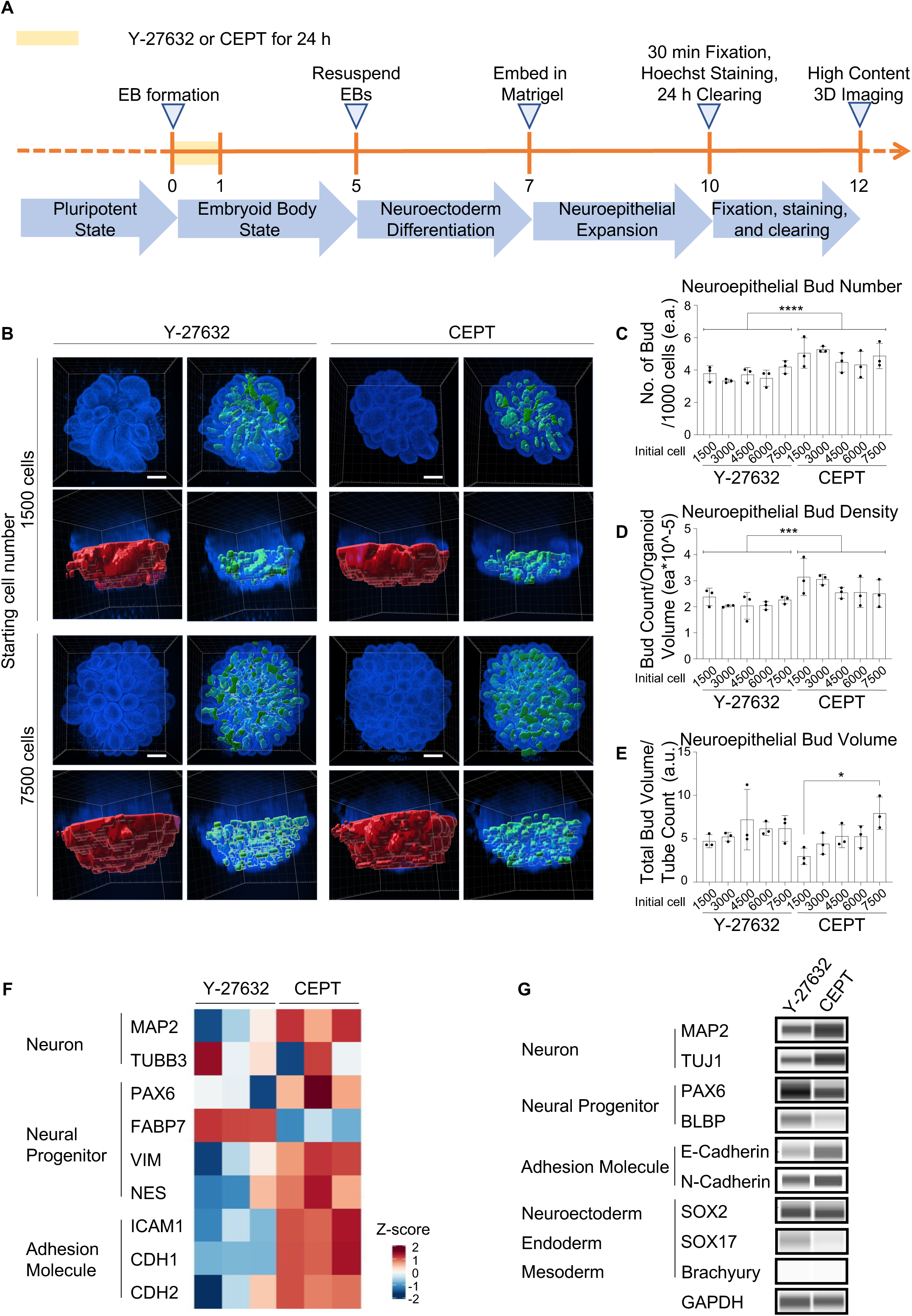
Analysis of early-stage brain organoids. (A) Overview of protocol for organoid generation and high-content 3D imaging. (B) Representative 3D rendered images of brain organoids generated either with Y-27632 or CEPT. Cell nuclei were stained with Hoechst. The region masked in red color represents the identified organoid volume and neuroepithelial buds (blue) were used for post-segmentation analysis. (C-E) Analysis of number, density, and volume of neuroepithelial buds in whole organoids based on segmentation analysis of Hoechst-labeled nuclei. (F) Bulk RNA-Seq and heatmap analysis of genes expressed by organoids (day 12) after treatment with Y-27632 or CEPT. (G) Western blot comparing expression of various markers in organoids generated with Y-27632 or CEPT. Note that neuronal markers MAP2 and TUJ1 are expressed at higher levels in the CEPT group and neural progenitor markers PAX6 and FABP7 are higher in the Y-27632 sample. See also the absence of the endoderm marker SOX17 after CEPT treatment. Scale bars, 300 µm.

Next, we performed RNA sequencing (RNA-seq) to examine early-stage organoid differentiation (day 12). Heatmap analysis of cell-type-specific genes (e.g., neural progenitors and neurons) and cell adhesion molecules involved in neuronal differentiation (Scuderi et al., 2021) showed marked differences between organoids generated with Y-27632 or CEPT (**Figure 3F**). These findings suggest that CEPT-generated organoids were relatively more differentiated and showed higher expression of neuronal-associated genes and cell adhesion molecules at day 12. To further corroborate these observations, early-stage organoids (day 12) were subjected to Western blot analysis. CEPT-treated organoids expressed higher levels of the neuronal marker MAP2 and cell adhesion molecules, along with lower levels of the neural progenitor markers PAX6 and FABP7 (also known as BLBP)(**Figure 3G**). SOX2 expression was at similar levels in both groups and the endoderm marker SOX17 was expressed in the Y-27632 group, while it was absent in CEPT-generated organoids. The mesoderm marker brachyury was not detected in either group. Together, these findings show that CEPT promotes neural differentiation in early-stage brain organoids when compared to Y-27632.

### CEPT treatment improves organoid architecture and *in vivo*-like differentiation

To investigate how early exposure to CEPT or Y-27632 may influence organoid biology at later stages, we cultured cerebral organoids for two months and performed several comparisons using different methods. Hematoxylin-eosin (H&E) staining showed the presence of compact neural rosette-like structures in CEPT-treated organoids, whereas Y-27632 exposure generated organoids with more variable morphologies (**Figure 4A**). These neural tube-like structures, which were generated more robustly with CEPT versus Y-27632, expressed the forebrain and neural progenitor markers FOXG1 and SOX2 (**Figures 4B and 4C**). As expected, neural tube-like structures were surrounded by neuronal cells expressing MAP2. Western blot analysis confirmed higher protein levels of markers expressed by neural progenitors (FOXG1, EN2, TBR2, and SOX2), neuroblasts (DCX), and more mature neurons (MAP2) (**Figure 4D**).

**Figure 4.**
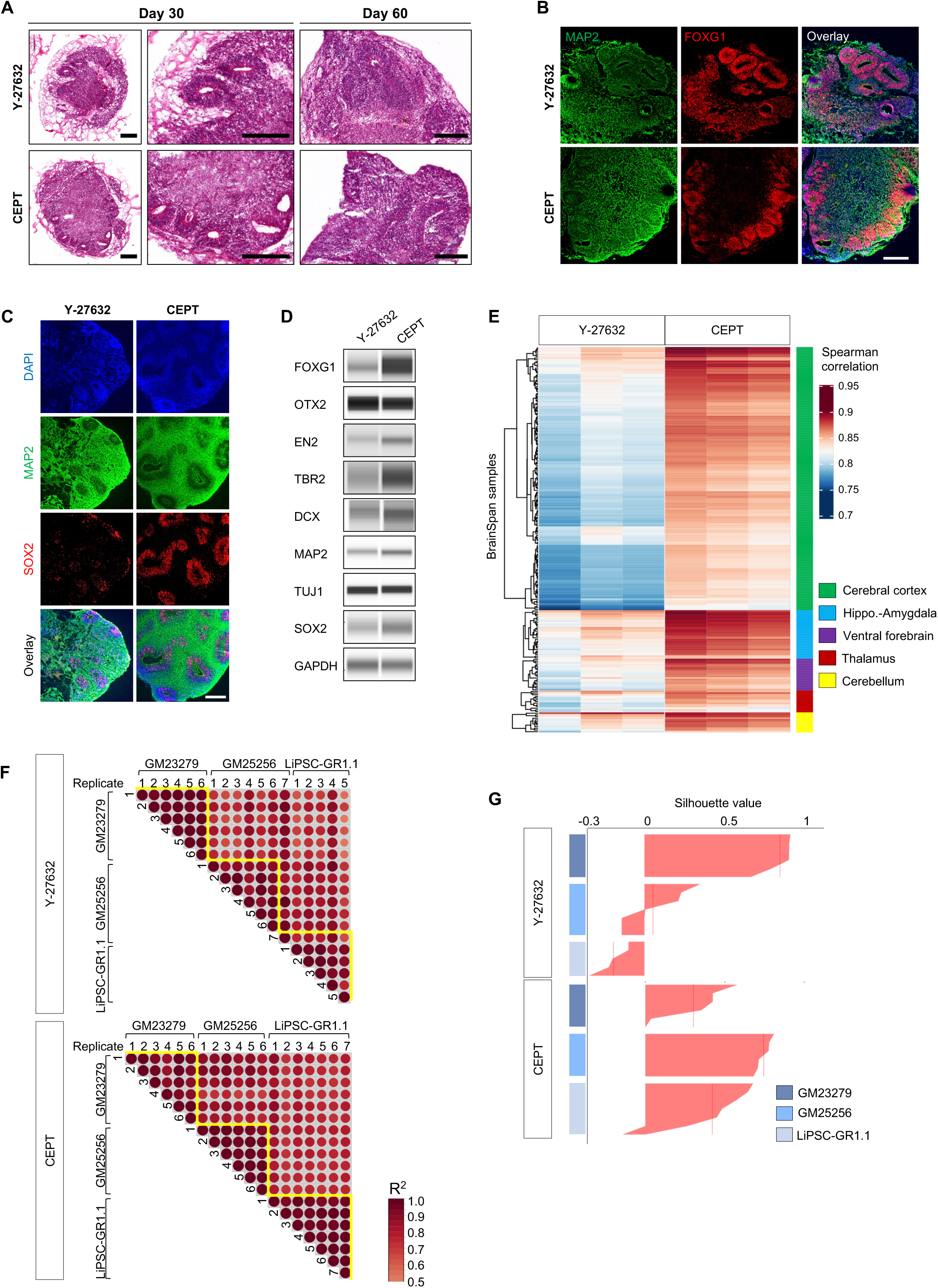
Neural architecture, comparison with brain tissues, and organoid reproducibility. (A) H&E staining of representative organoids at day 30 and day 60 derived with either Y-27632 or CEPT treatment. (B) Immunostaining of sectioned organoids at day 30 for forebrain marker FOXG1 and neuronal marker MAP2. Note the homogeneous formation of neural tube-like regions after CEPT treatment. (C) Immunostaining of MAP2-labeled neurons surrounding neural tube-like structures expressing SOX2 (day 60). Note that organoid anatomy is more consistent in the CEPT-generated example. (D) Western blot analysis of cell type- and brain-region-specific markers show differences between both groups (day 36). (E) Heatmap analysis (bulk RNA-Seq) and Spearman correlation coefficients reveal differences between organoids generated with Y-27632 and CEPT (day 60) and indicate similarity to the human cortex (Allen BrainSpan). (F) Pearson correlogram based on whole mRNA transcriptomes of individual organoids generated within the same and across different hiPSC lines at day 36. R^2^ values were computed with 95% confidence intervals. Deeper color represents a higher correlation (R^2^). (G) Silhouette plot based on transcriptomic analysis of individual organoids. Each bar represents single organoids derived from three cell lines. The silhouette value represents similarity within each cell line compared to the next most similar cell lines. Positive values indicate that the organoid is closer to other organoids within its cluster. The vertical line indicates the average silhouette value of each cluster. CEPT-generated organoids showed a closer average silhouette value compared to Y-27632 generated organoids. Scale bars, (A) 300 µm, (B) 300 µm, (C) 200 µm.

Next, we performed bulk RNA-seq analysis of organoids that were cultured for 2 months after 24-hour treatment with Y-27632 or CEPT. We compared the whole transcriptomes of these organoids to samples of the developing human cortex (8-9 weeks post-conception), which are part of the Allen BrainSpan Atlas (Miller et al., 2014). Indeed, this unbiased analysis using Spearman correlation revealed that the overall molecular signature of CEPT-generated organoids was more similar to early human brain tissue in contrast to Y-27632-generated organoids (**Figure 4E**).

Furthermore, we asked if CEPT may have beneficial effects on inter-organoid reproducibility and compared the transcriptomes of single organoids, which were generated from three different iPSC lines. Individual organoids (day 36) generated in parallel using Y-27632 or CEPT were processed for RNA-seq analysis. Again, we applied coefficient of determination (R^2^) analysis to measure similarity across the single cerebral organoids generated from the same iPSC line. We found substantial organoid-to-organoid variability after Y-27632 treatment as indicated by the low value in the correlation plot (**Figure 4F**) and the high distribution of the average silhouette value in the silhouette plot (**Figure 4G**). In contrast, CEPT-generated individual organoids displayed overall higher R^2^ values, indicating higher similarity. As shown in the silhouette plot, the comparison across cell lines also confirmed that CEPT reduces organoid-to-organoid variability (**Figures 4F and 4G**).

### Improved neuronal differentiation at single-cell resolution

To gain single-cell resolution, we performed single-cell RNA-seq (10X Genomics Chromium) for organoids generated with CEPT or Y-27632. We analyzed the transcriptomes of 8,952 cells (4,280 cells for the Y-27632 group; 4,672 cells for the CEPT group). Dimensionality reduction by principal component analysis (PCA) showed distinct global expression profiles for each group (**Figure 5A**). Top differentially expressed genes included VIM, NEUROD1, NEUROD2, NEUROD6, NFIB, DCX, MAP2, and NCAM1 (**Figure 5A**). Gene Ontology analysis based on the top 200 differentially expressed genes revealed that CEPT-generated organoids scored higher than Y-27632 organoids for neural-specific categories, including regulation of neurogenesis, neural development, axo-dendritic transport, glutamatergic synapse, and others (**Figure 5B**). We performed additional gene enrichment analysis using the ARCHS4 database (Lachmann et al., 2018), which allows unbiased comparison to 84,863 human transcriptomes. Again, when the top 200 differentially expressed genes comparing Y-27632 vs CEPT organoids were submitted, the top-hit categories indicated improved neural differentiation in CEPT organoids, whereas organoids generated using Y-27632 scored highest in non-neural categories such as “kidney” and “blood dendritic cells” (**Figure 5C**).

**Figure 5.**
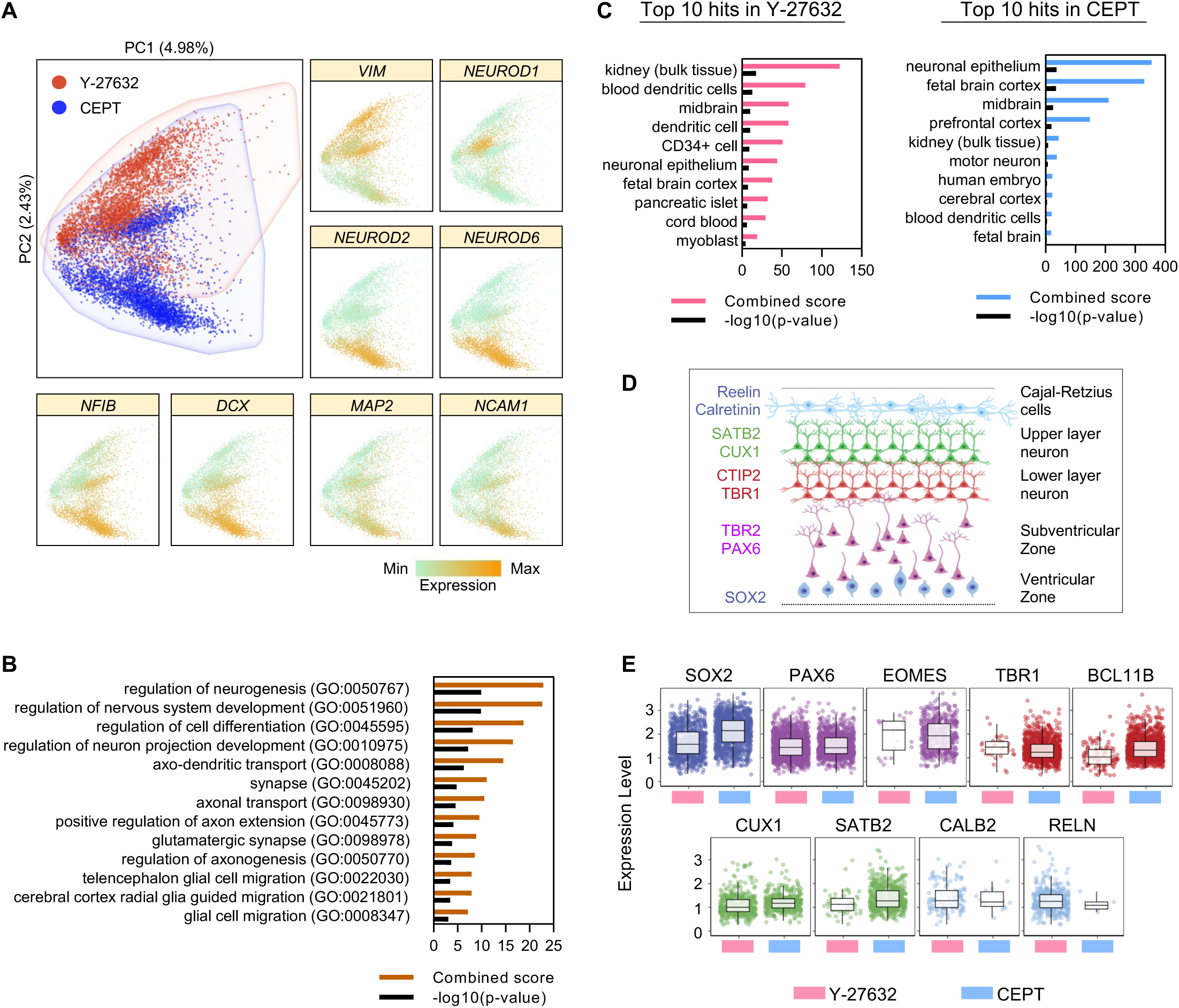
Single-cell RNA-seq of brain organoids generated with Y-27632 or CEPT. (A) Principal component analysis of organoids (day 72) shows distinct transcriptome profiles. Looking at the most strongly expressed genes, VIM and NEUROD1 were enriched in the Y-27632 organoids. Genes strongly expressed in CEPT organoids include NEUROD2, NEUROD6, NFIB, DCX MAP2, and NCAM1. (B) Gene ontology analysis of top 200 differentially expressed genes show that CEPT organoids score higher for neural and neuronal biological processes as compared to Y-27632. (C) EnrichR analysis comparing the transcriptomes of organoids generated with either Y-27632 or CEPT to human tissue samples in the ARCH4 database. CEPT organoids score higher for neural tissues than Y-27632. (D) Schematic of the cortical layers and genes expressed by different cell types. (E) Box plots (zero-value removed) showing relative gene expression of various cortical layers in organoids generated with Y-27632 or CEPT.

During brain development, the cortical layers are formed by migratory neuroblasts that migrate in an inside-out pattern from the subventricular zone toward the outer layers and the pial surface, whereby later-born neurons migrate through earlier formed cortical layers (Rowitch and Kriegstein, 2010). Using our single-cell RNA-seq dataset, we focused on analyzing cell-type-specific genes that are important for cortical layer formation (**Figure 5D**). Direct comparison of organoids revealed that CEPT treatment resulted in overall improved cortical development as indicated by higher and more consistent expression levels of various genes representing neural progenitors (SOX2, PAX6, EOMES/TBR2), as well as lower (TBR1, BCL11B/CTIP2) and upper layer neurons (CUX1, SATB2) (**Figure 5E**). Interestingly, only CALB2 and RELN, markers of Cajal-Retzius cells, were more abundant in the Y-27632 group. Given the transient role of Cajal-Retzius cells, this may indicate that CEPT organoids were relatively more mature. Lastly, we used immunohistochemistry and confirmed proper cortical layer formation (marked by expression of DCX, TBR1, TBR2, SATB2, and CTIP2)(**Figure S2**) in CEPT-generated organoids (day 60). Taken together, CEPT treatment during the first 24 hours of cell aggregation had long-lasting effects and improved neuronal differentiation of organoids cultured for over 2 months.

### Single-cell analysis confirms the superiority of CEPT organoids

To characterize the cell-type diversity in CEPT organoids relative to Y-27632 organoids, the cell populations were analyzed by unsupervised clustering algorithms based on global gene expression patterns and visualized on a two-dimensional t-distributed stochastic neighbor embedding (tSNE) plot (**Figure 6A**). t-SNE dimensionality reduction revealed seven distinct clusters where each cluster showed high differential gene expression compared to the rest, categorizing cell populations from CEPT organoids into three clusters and Y-27632 into four clusters. Unbiased cell population identification was performed based on a comparison of the expressed transcriptomes with a curated list of genes (**Supplementary Table S1**) for neural progenitors, radial glia, cortical neuron, and upper and lower cortical neurons based on a previous report (Trujillo et al., 2019). The data demonstrated that CEPT clusters (clusters 1-3) showed higher enrichment for molecular signatures representing cell types of developing cortex as compared to Y-27632 clusters (clusters 4-7)(**Figure 6A**). Gene set enrichment analysis based on the top 100 differentially expressed genes in each cluster showed significantly enriched neuronal development biological pathways in each cluster except for cluster 3 in CEPT and cluster 7 in Y-27632 (**Figures 6B and 6C; Supplementary Table S2 for full list of genes in Figure 6B**). While only a small fraction of cells in CEPT-generated organoids were non-neural (cluster 3), the non-neural cluster was larger in organoids generated using Y-27632 (cluster 7). Moreover, CEPT-generated organoids showed higher expression of several genes involved in neuronal function (SLC17A17, GRIN2B, KCNJ6, NR4A2, GABBR1, and GABBR2)(**Figure 6D**).

**Figure 6.**
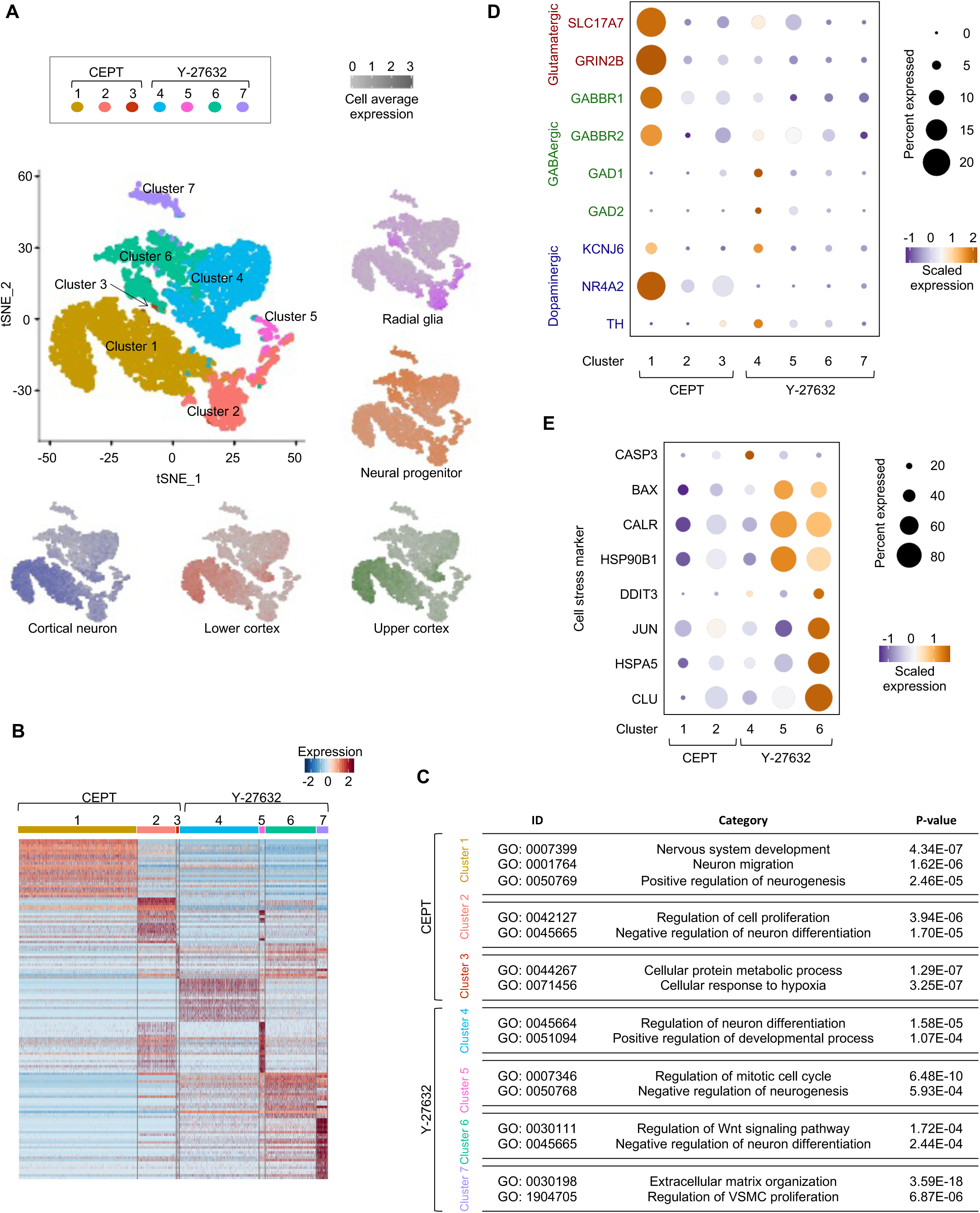
Sub-cluster analysis of organoids and stress markers. (A) UMAP plot for cell populations identified in organoids generated with Y-27632 or CEPT. Average expression of established cell type-specific markers. (B) Heatmap showing upregulated genes with a two-fold difference in each cluster. Each column represents a single cell clustered into unique sub-clusters based on similar gene expression profile and each row represents an individual gene. See Supplementary Table S2 for the complete gene list. Red indicates maximum gene expression, while blue indicates low or no expression in log-normalized UMI counts. (C) Gene ontology terms in relation to each cell cluster depicted in the heatmap in Figure 6B. (D) Dot plot showing expression levels for various neuronal markers and receptors in organoids generated with Y-27632 or CEPT. (E) Neuronal sub-clusters in CEPT-generated organoids indicate lower cell stress-related transcriptomics compared to those in Y-27632.

Cell stress-related markers were also found to be significantly higher in neuronal clusters derived with Y-27632 compared to CEPT. We performed Gene Ontology enrichment analysis using a curated list related to cell stress-related biological pathways (**Supplementary Table S3**) based on genes that were differentially expressed in Y-27632 versus CEPT-generated organoids. We found that the ER stress response was statistically higher in Y-27632 compared to CEPT-generated organoids, with the lowest expression of cell stress-related markers found in CEPT-generated neuronal clusters (**Figures 6E, S3A, and S3B**). These marked differences in both groups are important because activated cell stress pathways impede with differentiation in cortical organoids (Bhaduri et al., 2020).

### CEPT improves other neural and non-neural organoid models

To evaluate whether CEPT might be universally beneficial, we tested another brain organoid protocol (Velasco et al., 2019) and two commercially available kit-based methods (STEMCELL Technologies) that generate intestinal and kidney organoids based on previous reports (Freedman et al., 2015; Spence et al., 2011). First, we generated brain organoids in parallel using 20 µM Y-27632 (Velasco et al., 2019) or CEPT (**Figure S4A**). Organoids of both treatment groups were cultured until day 35 and then processed for immunohistochemical analysis. CEPT-generated organoids showed improved anatomical organization with prominent neural tube-like structures expressing PAX6 surrounded by neuronal cells expressing MAP2. In contrast, organoids generated by using Y-27632 were more variable and more challenging to analyze histologically (**Figure S4B**).

Next, we generated intestinal organoids from two hPSC lines (WA09 and LiPSC-GR1.1) treated with Y-27632 following the manufacturer’s instructions (STEMCELL Technologies) or treated with CEPT (**Figure 7A**). At day 28, organoids were fixed and processed for immunohistochemical analysis using specific antibodies against mucin 2 (a marker for goblet cells), CDX2 (a marker for intestinal epithelium), ZO-1 (a marker for lumen formation), and the proliferation marker Ki67. This comparison showed that CEPT treatment generated improved intestinal organoids with tissue-specific architecture, including polarized epithelial cells surrounding lumen-like structures (**Figure 7B**). In comparison to Y-27632, intestinal organoids were generally larger and displayed more complex morphologies.

**Figure 7.**
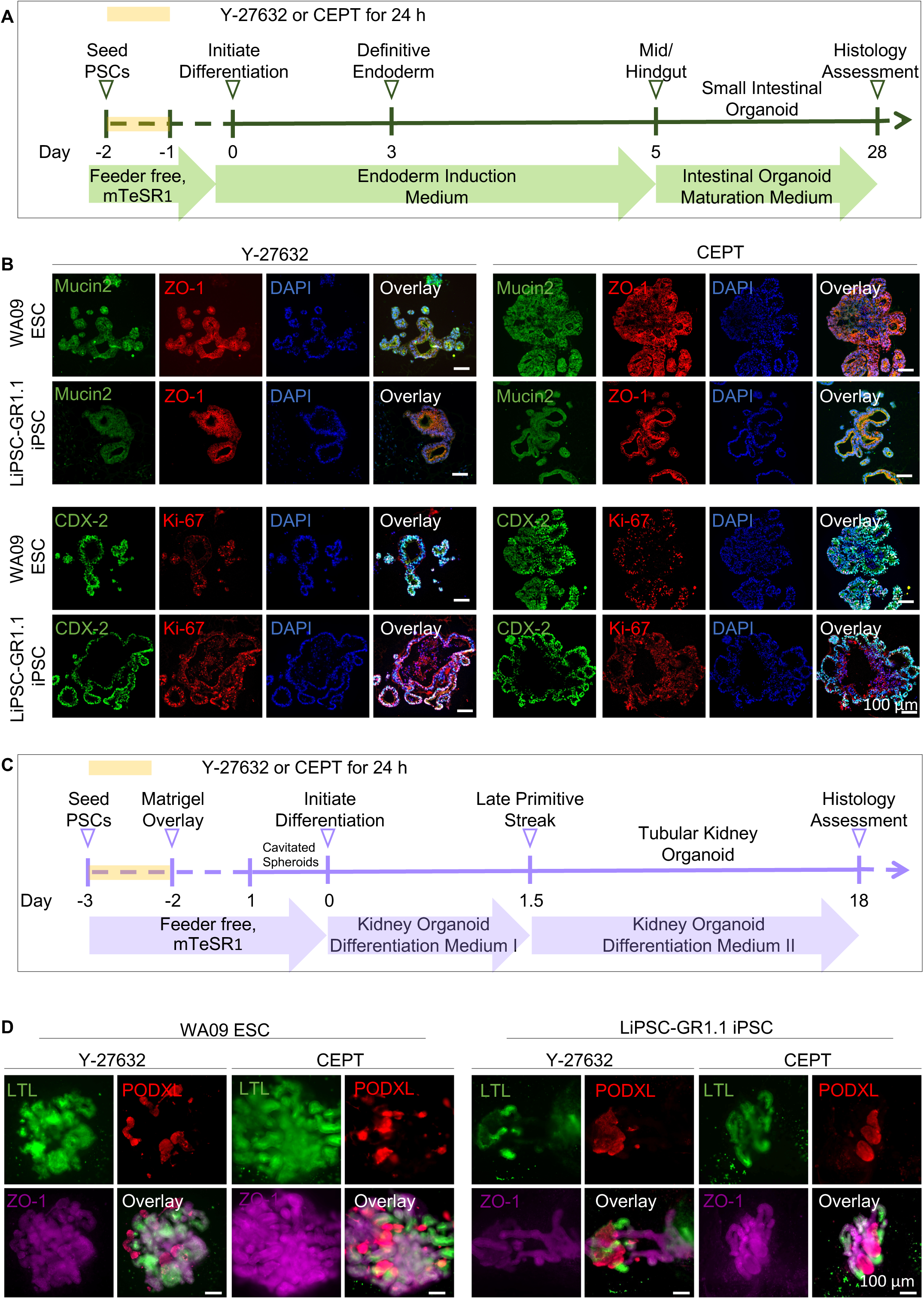
CEPT cocktail improves generation of intestinal and kidney organoids. (A) Overview of 28-day protocol generating intestinal organoids. (B) Immunohistochemical analysis of intestinal organoids using various relevant markers. Organoids were generated in parallel from two cell lines (WA09 and LiPSC-GR1.1). (C) Overview of 18-day protocol generating kidney organoids. (D) Immunohistochemical analysis of kidney organoids using specific markers. Organoids were generated in parallel from two cell lines (WA09 and LiPSC-GR1.1). Scale bars, 100 µm.

Kidney organoids were generated using Y-27632 according to the manufacturer’s recommendation or using CEPT treatment (**Figure 7C**). At day 18, organoids from the two hPSC lines were fixed and processed for whole-mount immunohistochemistry. Nephron segments were stained using proximal tubule marker LTL (lotus tetragonolobus lectin), podocyte marker PODXL (podocalyxin), and epithelial marker (ZO-1) (**Figure 7D**). Immunostainings and phase-contrast microscopy showed clear differences between organoids generated with Y-27632 or CEPT (**Figures 7D** and **S5**). Kidney organoids generated using CEPT were larger and displayed more complex morphologies as compared to Y-27632 organoids.

### Discussion

Cell aggregation and self-organization have been used for over a century to study the simple metazoan *Hydra* (Layer et al., 2002). More recently, these tissue engineering methods have been employed to generate various stem cell-based 3D models such as EBs, neurospheres, and organoids from different developmental lineages (Eiraku et al., 2008; Keller, 1995; Lancaster et al., 2013; Quadrato et al., 2017; Reynolds and Weiss, 1992; Sato et al., 2009; Singec et al., 2006; Tristan et al., 2021; Tsankov et al., 2015). Culturing hPSCs is particularly challenging as these cells are inherently sensitive to environmental perturbations and require special conditions, cell culture media, and reagents. In addition, organoids ectopically activate cellular stress pathways that compromise cell type specification (Bhaduri et al., 2020) and other limitations of organoids have been described elsewhere (Andrews and Kriegstein, 2022). We recently reported the development of the CEPT small molecule cocktail, which dramatically improves cell viability by preventing multiple stress mechanisms and DNA damage (Chen et al., 2021). Here we extend these observations and provide detailed analyses of EBs and four different organoid models that benefit from CEPT treatment. We consistently observed that CEPT treatment was superior to Y-27632. CEPT improves cell survival, prevents cellular stress, leads to optimal cell aggregation, reduces experimental variability, results in overall improved cell differentiation and tissue-specific 3D architecture. The four-part CEPT cocktail is also more cost-efficient than 10 µM Y-27632 and other reagents (Chen et al., 2021, Tristan et al., 2021). Hence, the use of CEPT promotes standardization, optimal biological outcome, and should help with utilizing next-generation EB and organoid models suitable for translational applications and drug discovery.

### Experimental Procedures

Detailed descriptions of experimental procedures are provided in the supplemental information.

### Chemical compounds

ROCK inhibitor Y-27632 (Tocris, 10 µM, unless otherwise stated) and CEPT cocktail components (Chroman 1, MedChemExpress, 50 nM; Emricasan, Selleckchem 5 µM; Polyamine, Sigma Aldrich, 1:1000; trans-ISRIB, Tocris, 7 µM) were prepared following the manufacturer’s recommendations.

### Cell culture and brain organoid formation

All hESC (WA09) and hiPSC (LiPSC-GR1.1, GM23279, and GM25256) lines were maintained under feeder-free conditions in mTeSR medium (STEMCELL Technologies) and VTN-N-coated (Thermo Fisher Scientific) plates as described in the supplemental information. Cerebral organoids were generated using a commercial kit (STEMdiff™ cerebral organoid kit, STEMCELL Technologies) or a previously published protocol (Velasco et al., 2019). Protocols were summarized in Figure 1A and S4A with details described in the supplemental information.

### EB viability analysis

Phase-contrast images of EBs were obtained with an Incucyte Zoom Live Cell Analysis System (Sartorius). The CellTiter-Glo 3D assay (Promega) was carried out following the manufacturer’s instructions to quantify cell viability. Luminescence signal was read using the PHERAstar FXS microplate reader (BMG LABTECH). EB size was measured using the Celigo Imaging Cytometer (Nexcelom Biosciences). Live and dead cells were stained using a two-color fluorescence live/dead assay kit (Thermo Fisher Scientific, 1:2,000) and imaged with an DMi8 microscope (Leica) or an Opera Phenix high-content microscope (PerkinElmer) using appropriate filters. During each assessment, single EBs were cultured in 96-well ULA round-bottom plates (Corning).

### Western Blot

Western blot analyses were performed using the Wes automated western blotting system (Protein Simple) following the manufacturer’s instruction. Information on primary and secondary antibodies are provided in Supplementary Table S4.

### RASL-seq analysis of single EBs

hESCs (WA09) were dissociated and cultured at a density of 20,000 cells per well in 96-well ULA round-bottom plates (Corning) in E6 medium (Thermo Fisher Scientific) for 7 days. As positive controls for lineage-specific gene expression, hESCs were differentiated into ectoderm, mesoderm, and endoderm with details described in the Supplemental Information. Single EBs were harvested, lysed in TCL lysis buffer (Qiagen), PCR amplified, and purified. Purified samples were sequenced on an Illumina NextSeq 550 using custom sequencing primers. Protocol overview is presented in Figure 2A and method details are provided in Supplemental Information.

### High-content imaging and analysis of organoids

Cerebral organoids of varying starting cell numbers (1,500-7,500 cells per organoid) were generated following the kit-based protocol (STEMCELL Technologies) and fixed with 4% PFA at day 10. The fixed organoids were stained with Hoechst 33342 (Thermo Fisher Scientific), cleared with clearing solution, and imaged on the Opera Phenix high-content imaging microscope (PerkinElmer). A customized image analysis script was used for automated measurement of sphere features. Protocol overview is presented in Figure 3A and method details are provided in Supplemental Information.

### Bulk RNA-Sequencing

Three organoids were pooled to prepare each sample (three samples per group). RNA was extracted using the RNeasy Mini Kit (Qiagen). For reproducibility studies, RNA was extracted from single organoids. Extracted RNA was prepped for RNA-seq libraries with the TruSeq Stranded mRNA Library Prep Kit (Illumina) and sequenced using the Illumina NextSeq 550 system according to the manufacturer’s protocol. Methods describing bioinformatic analysis, including correlation analysis, can be found in Supplemental Information.

### Histological analysis

Organoids were fixed in 4% PFA, washed with PBS, immersed in 30% sucrose overnight, embedded in O.C.T. compound (Fisher Scientific), cut into 20 µm sections, and mounted on microscope slides. For histological analysis, sections were stained with H&E, dehydrated, and mounted with Permount mounting medium (Fisher Scientific). For immunohistochemical analysis, sections were permeabilized, blocked, stained with primary and secondary antibodies, and mounted with ProLong Glass Antifade Mountant with NucBlue Stain (Thermo Fisher Scientific). Images were taken with the Zeiss LSM 710 confocal microscope. Detailed information on primary and secondary antibodies is provided in Supplementary Table S4.

### Single-cell RNA Sequencing library preparation and analysis

Organoids at day 72 of culture were dissociated into single cells using the Embryoid Body Dissociation kit (Miltenyi Biotec) and gentle MACS Dissociator (Miltenyi Biotech) following the manufacturer’s protocol. The strained cell suspension was loaded on a Chromium Controller (10X Genomics) to generate single-cell gel bead-in-emulsions (GEM) and barcoding. The library was sequenced on an Illumina NextSeq 550. Details of the library preparation and analysis procedure are described in the Supplemental Information.

### Intestinal and kidney organoids formation

Intestinal and kidney organoids were generated using commercial kits (STEMCELL Technologies) following the manufacturer’s protocol. Protocol overviews are shown in Figure 7A and 7C and method details are described in the Supplemental Information.

### Data and code availability

Bulk and single-cell RNA-seq data generated in this study can be found in the NCBI SRA under the Bioproject PRJNA815659. Processed RASL-Seq data files can be accessed at GEO GSE198575 (raw data: PRJNA816058).

The umbrella SRA Bioproject is PRJNA816454. Analysis code is available at https://github.com/cemalley/Ryu_cerebral_organoids.

## SUPPLEMENTAL INFORMATION

Supplemental information can be found online at https://

## AUTHOR CONTRIBUTIONS

S.R. and I.S. conceived the study and experiments. S.R., P.C., C.A.T., V.M.J., T.D., H.H., T.C.V. performed experiments. S.R., C.W., C.A.T., B.E., V.M.J., T.D., J.S., H.H., J.B., M.H., A.S., T.C.V., and I.S. contributed to data analysis and discussions. S.R. and I.S. wrote the manuscript.

### Conflict of Interests

A.S. and I.S. are co-inventors on a U.S. Department of Health and Human Services patent application covering the CEPT cocktail and its use.

## Acknowledgements

We are grateful for the support from the Regenerative Medicine Program (RMP) of the NIH Common Fund, NIH HEAL Initiative, and in part by the intramural research program of the National Center for Advancing Translational Sciences (NCATS), NIH. We thank Hannah Baskir for critical reading of the manuscript. The funders had no role in study design, data collection and analysis, decision to publish, or preparation of the manuscript. Figure 5D was created with BioRender.com.

## SUPPLEMENTAL MOVIE

**Movie S1: Brain organoid generated with CEPT**

The exemplary movie demonstrates the 3D morphology of a CEPT-generated brain organoid at the end of the neuroepithelial expansion stage (day 10). After optical clearing, confocal images were taken and digitally analyzed to quantify organoid volume, neural tube numbers, and single cell numbers. See also Figures 3A-3D.

