## Supplemental Information for "Enhancing the Fitness of Embryoid Bodies and Organoids by Chemical Cytoprotection"

**Figure S1**

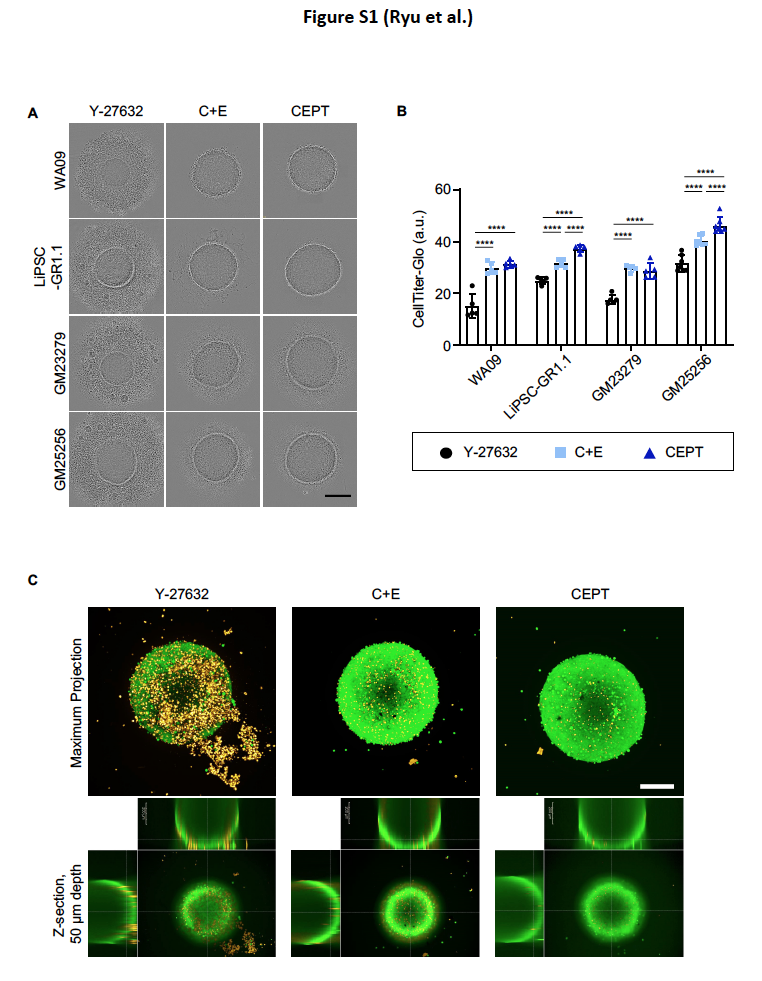

**Figure S1: Improved cell viability and EB formation with CEPT**

(A) Phase-contrast images of EBs generated from hESCs (WA09) and three different iPSC lines (LiPSC-GR1.1, GM23279, GM25256).
(B) Quantifications of cell survival in EBs with the CellTiter-Glo assay measuring ATP levels.
(C) Confocal images of EBs projected at maximum intensity (z-scan of 50 µm from the surface) to visualize live and dead cells that are inside and outside of EBs.

Images were collected from EBs at day 3 (A-B) and day 5 (C).

**Figure S2**

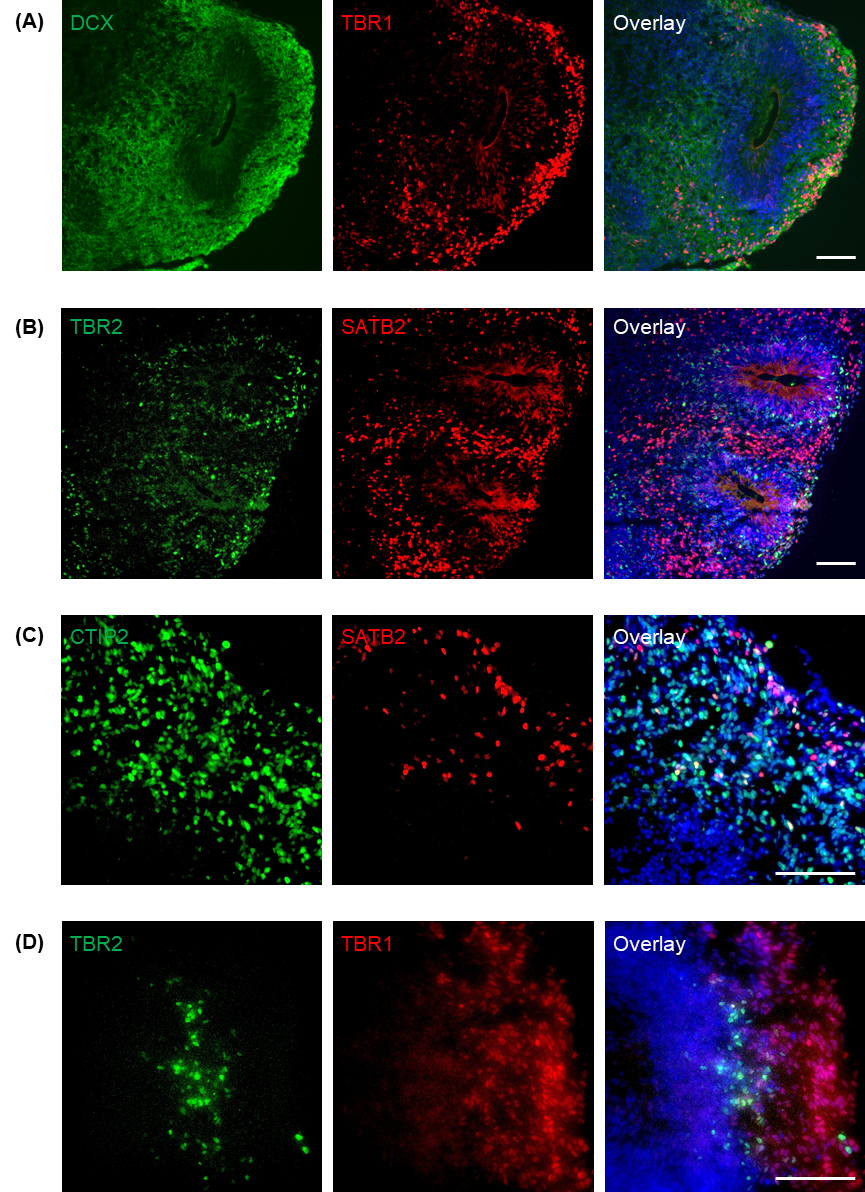

**Figure S2: Cortical layers in brain organoids generated with CEPT**

Immunohistochemical analysis of brain organoids (day 60) to identify cortical neurons.

(A) Neuroblast marker DCX and post-mitotic neuron marker TBR1.

(B) Deep-layer cortical neuron marker TBR2 and upper-layer cortical neuron maker SATB2.

(C) Deep-layer cortical neuron marker CTIP2 and upper-layer cortical neuron marker SATB2.

(D) Neurons in different cortical layers express TBR1 and TBR2.

The data demonstrate proper inside-out migration and positioning. Note that neurons in the deep layers express TBR2 and CTIP2, and cells closer to the organoid surface express TBR1 and SATB2. See also schematic in Figure 5D.

Scale bars, 100 µm.

**Figure S3**

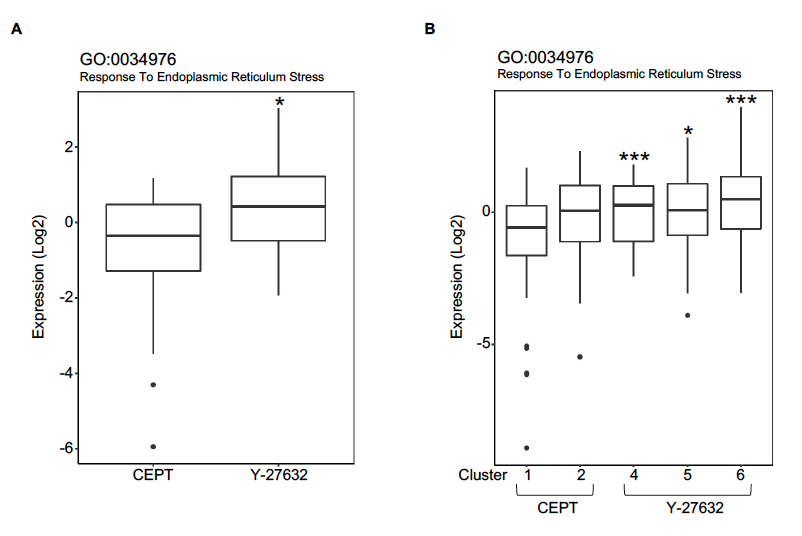

**Figure S3: Gene set enrichment analysis of cellular stress in brain organoids**

(A) Gene ontology analysis of single-cell RNA-seq data showing elevated ER stress genes in organoids generated with Y-27632 versus CEPT. Organoids from both groups were analyzed at day 72.

(B) Sub-cluster analysis also indicates a trend for higher levels of ER stress in Y-27632 versus CEPT. Cluster 1 represents the neuronal population in CEPT organoids. P value was calculated in comparison to Cluster 1. P values represent *p<0.05, ***p<0.001.

**Figure S4**

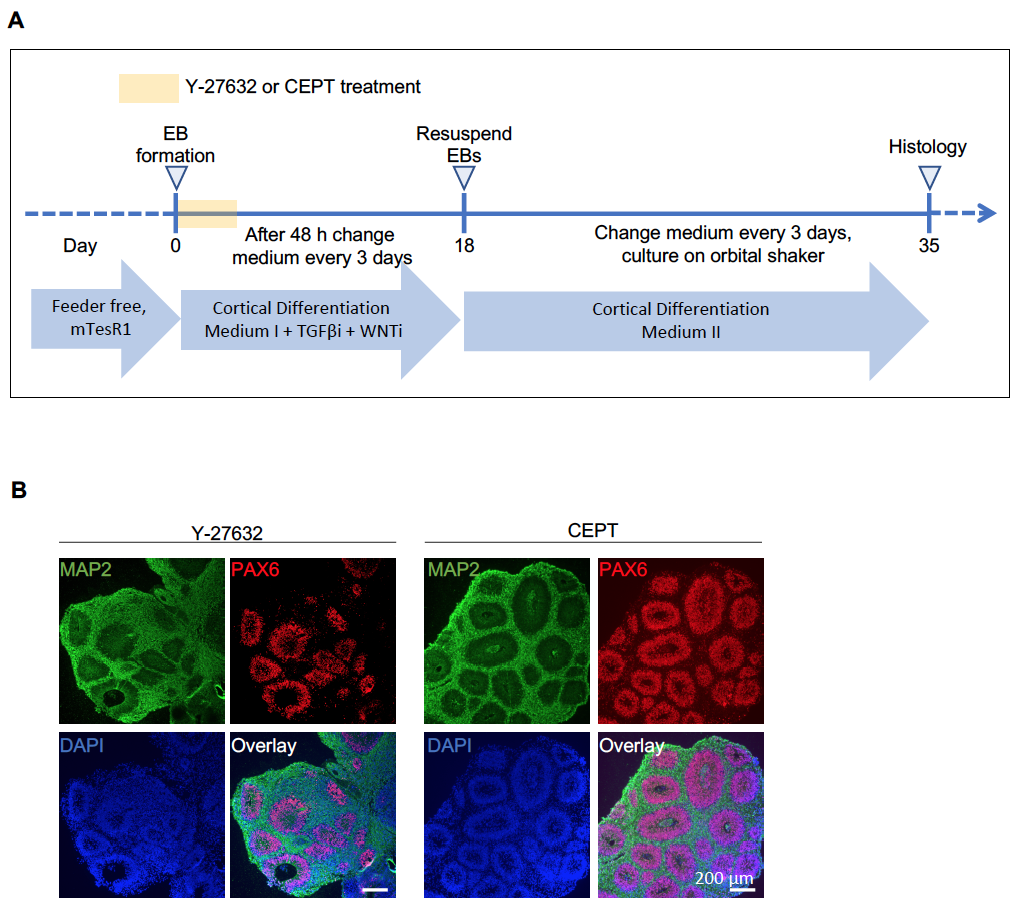

**Figure S4. Reproducible improvement of organoid architecture by CEPT**

(A, B) Brain organoids were generated using a published protocol (Velasco et al., 2019) and analyzed at day 35. Replacing Y-27632 with the CEPT cocktail during cell aggregation resulted in improved neural tube-like structures expressing PAX6. Neuronal cells were visualized by MAP2 immunostaining.

Scale bars, 200 µm.

**Figure S5**

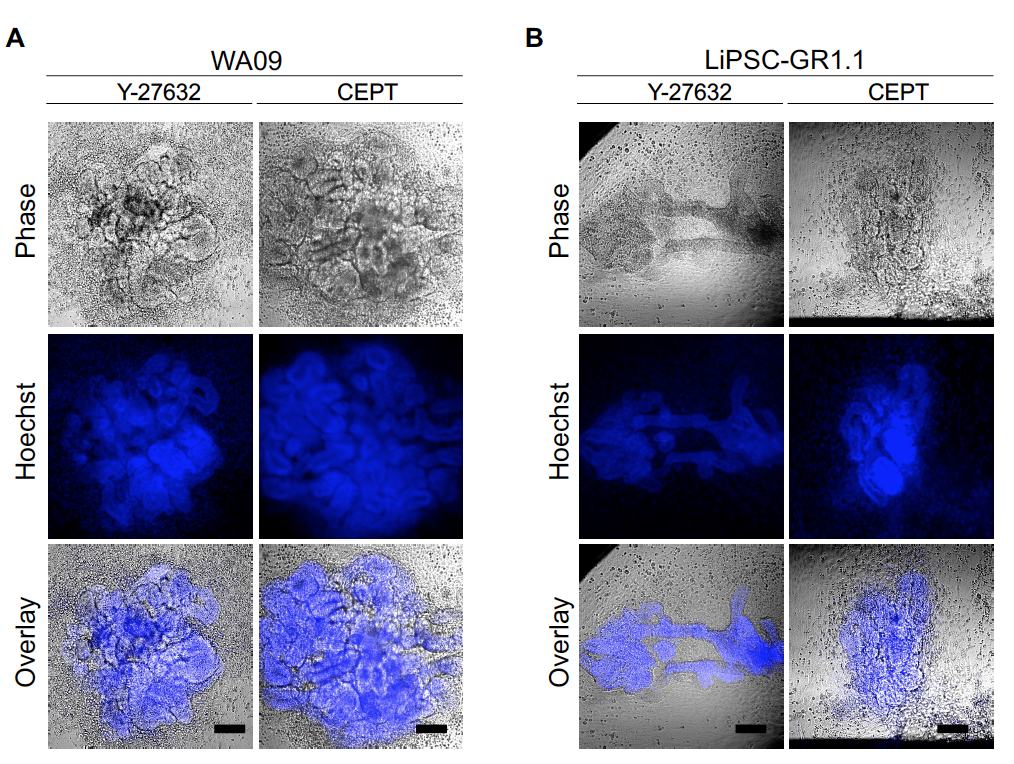

**Figure S5. Microscopic overview of kidney organoids**

Phase-contrast images overlayed with fluorescent images of kidney organoids generated using a kit-based protocol (STEMCELL Technologies). Note that organoid size and morphology are affected by Y-27632 or CEPT treatment as reproduced using two cell lines, (A) WA09 ESC and (B) LiPSC-GR1.1 iPSC.

Scale bars, 100 µm.

**Table S1. Curated list of neuronal genes**

| Category | Gene |
| --- | --- |
| Cortical neurons | SOX11 |
| Cortical neurons | NEUROD2 |
| Cortical neurons | SOX2 |
| Cortical neurons | GPM6A |
| Cortical neurons | SOX4 |
| Cortical neurons | MLLT11 |
| Cortical neurons | CCNI |
| Cortical neurons | SLA |
| Cortical neurons | MARCKSL1 |
| Cortical neurons | DCX |
| Cortical neurons | NES |
| Neural Progenitor | ANXA2 |
| Neural Progenitor | GYPC |
| Neural Progenitor | SPARC |
| Neural Progenitor | SDC2 |
| Neural Progenitor | CRABP2 |
| Neural Progenitor | NTRK2 |
| Neural Progenitor | CCND1 |
| Neural Progenitor | LGALS1 |
| Neural Progenitor | SERF2 |
| Neural Progenitor | MDK |
| Neural Progenitor | VGLL3 |
| Neural Progenitor | S100A13 |
| Neural Progenitor | PDLIM7 |
| Neural Progenitor | ANXA5 |
| Neural Progenitor | PRSS23 |
| Neural Progenitor | RPL41 |
| Neural Progenitor | NPC2 |
| Neural Progenitor | SEC11A |
| Neural Progenitor | PRDX6 |
| Neural Progenitor | TPM1 |
| Neural Progenitor | RHOC |
| Neural Progenitor | NEAT1 |
| Neural Progenitor | RPL12 |
| Neural Progenitor | RPL7A |
| Neural Progenitor | EEF1A1 |
| Neural Progenitor | RPL28 |
| Neural Progenitor | RPS6 |
| Neural Progenitor | RPL23A |
| Neural Progenitor | TIMP1 |
| Neural Progenitor | RPL8 |
| Neural Progenitor | METRN |
| Neural Progenitor | WLS |
| Neural Progenitor | RPL27A |
| Neural Progenitor | CTGF |
| Neural Progenitor | RCN1 |
| Neural Progenitor | PFN1 |
| Neural Progenitor | PMP22 |
| Neural Progenitor | ITGB8 |
| Neural Progenitor | SERPINH1 |
| Neural Progenitor | VIM |
| Neural Progenitor | NME4 |
| Neural Progenitor | RPS7 |
| Neural Progenitor | MYL12A |
| Neural Progenitor | RPS20 |
| Neural Progenitor | RPS2 |
| Neural Progenitor | RPLP1 |
| Neural Progenitor | RAB13 |
| Neural Progenitor | TUBB6 |
| Neural Progenitor | CRNDE |
| Neural Progenitor | TTYH1 |
| Neural Progenitor | RPL23 |
| Neural Progenitor | RPS19 |
| Neural Progenitor | RPL29 |
| Neural Progenitor | RPS14 |
| Neural Progenitor | RPL3 |
| Neural Progenitor | SLC25A6 |
| Neural Progenitor | SPATS2L |
| Neural Progenitor | QPRT |
| Neural Progenitor | RPL35 |
| Neural Progenitor | RPS18 |
| Neural Progenitor | CLIC1 |
| Neural Progenitor | RPS3 |
| Neural Progenitor | RPL10A |
| Neural Progenitor | RPS28 |
| Neural Progenitor | CD63 |
| Neural Progenitor | PDPN |
| Neural Progenitor | ACTG1 |
| Neural Progenitor | CCNG1 |
| Neural Progenitor | CD99 |
| Neural Progenitor | B2M |
| Neural Progenitor | CHCHD10 |
| Neural Progenitor | RPLP0 |
| Neural Progenitor | RPS27L |
| Neural Progenitor | COL1A2 |
| Neural Progenitor | PFN2 |
| Neural Progenitor | UBB |
| Neural Progenitor | RPL37 |
| Neural Progenitor | CRABP1 |
| Neural Progenitor | RPL7 |
| Neural Progenitor | FSTL1 |
| Neural Progenitor | RPL36 |
| Neural Progenitor | RPL19 |
| Neural Progenitor | FGFR1 |
| Neural Progenitor | ENO1 |
| Neural Progenitor | RPS15 |
| Neural Progenitor | MYL6 |
| Neural Progenitor | GSTP1 |
| Neural Progenitor | PODXL |
| Neural Progenitor | CNN3 |
| Neural Progenitor | GNG11 |
| Neural Progenitor | RPS4Y1 |
| Neural Progenitor | AHNAK |
| Neural Progenitor | CST3 |
| Neural Progenitor | RPS23 |
| Neural Progenitor | RPL13A |
| Radial Glia | SFRP1 |
| Radial Glia | SOX2 |
| Radial Glia | C1orf61 |
| Radial Glia | FABP7 |
| Radial Glia | SLC1A3 |
| Radial Glia | SYNE2 |
| Radial Glia | PAX6 |
| Radial Glia | HMGN3 |
| Radial Glia | ID4 |
| Radial Glia | MYO10 |
| Radial Glia | DBI |
| Radial Glia | PTN |
| Radial Glia | QKI |
| Radial Glia | LINC01158 |
| Radial Glia | ZFHX4 |
| Radial Glia | HES1 |
| Radial Glia | HMGB2 |
| Radial Glia | LHX2 |
| Lower cortex | SNAP25 |
| Lower cortex | GRIA2 |
| Lower cortex | CNTNAP2 |
| Lower cortex | CELF4 |
| Lower cortex | NSG2 |
| Lower cortex | SYT1 |
| Lower cortex | YWHAH |
| Lower cortex | SNCA |
| Lower cortex | BASP1 |
| Lower cortex | DOK6 |
| Lower cortex | RTN1 |
| Lower cortex | RUNX1T1 |
| Lower cortex | FAM49A |
| Lower cortex | MAP1B |
| Lower cortex | SYT4 |
| Lower cortex | B3GALT2 |
| Lower cortex | GABRB2 |
| Lower cortex | LMO3 |
| Lower cortex | SCG3 |
| Lower cortex | UCHL1 |
| Lower cortex | VAMP2 |
| Lower cortex | TMEM161B -AS1 |
| Lower cortex | LY6H |
| Lower cortex | MAPT |
| Lower cortex | CDKN2D |
| Lower cortex | RAB3A |
| Upper cortex | MEF2C |
| Upper cortex | STMN2 |
| Upper cortex | NSG2 |
| Upper cortex | ARPP21 |
| Upper cortex | STMN4 |
| Upper cortex | MAPT |
| Upper cortex | GRIN2B |
| Upper cortex | CALM1 |
| Upper cortex | NELL2 |
| Upper cortex | SCD5 |
| Upper cortex | SATB2 |
| Upper cortex | PKIA |
| Upper cortex | MAP1B |
| Upper cortex | INA |
| Upper cortex | STMN1 |
| Upper cortex | NEUROD6 |
| Upper cortex | VAMP2 |
| Upper cortex | DOK5 |
| Upper cortex | RASL11B |
| Upper cortex | SNCA |
| Upper cortex | R3HDM1 |
| Upper cortex | TTC9B |
| Upper cortex | RAC3 |
| Upper cortex | CXADR |
| Upper cortex | HN1 |
| Upper cortex | CAMK2B |
| Upper cortex | RTN1 |
| Upper cortex | CHL1 |
| Upper cortex | NSG1 |
| Upper cortex | TUBB2A |
| Upper cortex | GABBR2 |
| Upper cortex | RBFOX2 |
| Upper cortex | CRMP1 |
| Upper cortex | GAP43 |
| Upper cortex | UCHL1 |
| Upper cortex | CDKN2D |
| Upper cortex | NCAM1 |
| Upper cortex | MSRA |
| Upper cortex | GPR85 |
| Upper cortex | DAAM1 |

**Table S2. List of differentially expressed genes in cell clusters identified by single-cell RNA-seq analysis (day 72 organoids)**

| p_val | avg_logFC | pct.1 | pct.2 | p_val_adj | cluster | gene |
| --- | --- | --- | --- | --- | --- | --- |
| 0 | 2.79308833 | 0.938 | 0.098 | 0 | cluster1 | NEUROD6 |
| 0 | 2.05248285 | 0.856 | 0.091 | 0 | cluster1 | NEUROD2 |
| 0 | 1.77946201 | 0.933 | 0.493 | 0 | cluster1 | SOX11 |
| 0 | 1.50330742 | 0.986 | 0.814 | 0 | cluster1 | SOX4 |
| 0 | 1.47253038 | 0.901 | 0.369 | 0 | cluster1 | DCX |
| 0 | 1.44982571 | 0.914 | 0.422 | 0 | cluster1 | NFIB |
| 0 | 1.40268984 | 0.6 | 0.016 | 0 | cluster1 | SLA |
| 0 | 1.30627288 | 0.685 | 0.14 | 0 | cluster1 | BCL11A |
| 0 | 1.28895579 | 0.917 | 0.477 | 0 | cluster1 | GPM6A |
| 0 | 1.23937348 | 0.412 | 0.048 | 0 | cluster1 | MEF2C |
| 0 | 1.22895865 | 0.955 | 0.691 | 0 | cluster1 | STMN2 |
| 0 | 1.22621736 | 0.877 | 0.375 | 0 | cluster1 | RTN1 |
| 0 | 1.17589242 | 0.755 | 0.363 | 0 | cluster1 | CSRP2 |
| 0 | 1.15102301 | 0.6 | 0.126 | 0 | cluster1 | NELL2 |
| 0 | 1.10309116 | 0.82 | 0.348 | 0 | cluster1 | CRMP1 |
| 0 | 1.08903613 | 0.688 | 0.127 | 0 | cluster1 | FOXG1 |
| 0 | 1.06145477 | 0.755 | 0.421 | 0 | cluster1 | ID2 |
| 3.31E-184 | 1.03536496 | 0.291 | 0.07 | 1.10E-179 | cluster1 | LMO3 |
| 0 | 1.0199526 | 0.946 | 0.727 | 0 | cluster1 | MLLT11 |
| 0 | 1.01720032 | 0.507 | 0.047 | 0 | cluster1 | CAMKV |
| 0 | 1.0057951 | 0.672 | 0.172 | 0 | cluster1 | PPP2R2B |
| 0 | 1.00231164 | 0.949 | 0.728 | 0 | cluster1 | JPT1 |
| 0 | 1.00023046 | 0.507 | 0.072 | 0 | cluster1 | GRIA2 |
| 0 | 1.73431764 | 0.828 | 0.157 | 0 | cluster2 | SFRP1 |
| 0 | 1.72310132 | 0.861 | 0.124 | 0 | cluster2 | SOX2 |
| 0 | 1.46288089 | 0.8 | 0.213 | 0 | cluster2 | TTYH1 |
| 0 | 1.34463153 | 0.99 | 0.557 | 0 | cluster2 | C1orf61 |
| 0 | 1.32085618 | 0.992 | 0.709 | 0 | cluster2 | FABP7 |
| 4.50E-160 | 1.30209754 | 0.638 | 0.291 | 1.49E-155 | cluster2 | HMGB2 |
| 1.61E-289 | 1.29772835 | 0.404 | 0.06 | 5.32E-285 | cluster2 | PCLAF |
| 0 | 1.2372285 | 0.955 | 0.661 | 0 | cluster2 | FABP5 |
| 7.90E-195 | 1.2246625 | 0.434 | 0.109 | 2.61E-190 | cluster2 | PTTG1 |
| 1.25E-10 | 1.17850122 | 0.419 | 0.361 | 4.13E-06 | cluster2 | HIST1H4C |
| 0 | 1.12174966 | 0.521 | 0.019 | 0 | cluster2 | SOX3 |
| 5.06E-259 | 1.05383338 | 0.658 | 0.229 | 1.67E-254 | cluster2 | PEA15 |
| 0 | 1.03579857 | 0.599 | 0.089 | 0 | cluster2 | EMX2 |
| 0 | 1.0186682 | 0.614 | 0.116 | 0 | cluster2 | NKAIN3 |
| 3.14E-246 | 1.01257066 | 0.494 | 0.115 | 1.04E-241 | cluster2 | MT3 |
| 5.53E-72 | 1.00623917 | 0.558 | 0.331 | 1.83E-67 | cluster2 | HES6 |
| 3.61E-273 | 1.00272802 | 0.943 | 0.634 | 1.19E-268 | cluster2 | VIM |
| 4.61E-165 | 1.00181194 | 0.257 | 0.04 | 1.52E-160 | cluster2 | TOP2A |
| 0 | 1.00119856 | 0.766 | 0.244 | 0 | cluster2 | DOK5 |
| 2.97E-17 | 1.92945483 | 0.478 | 0.142 | 9.83E-13 | cluster3 | CRYAB |
| 9.43E-08 | 1.57132013 | 0.246 | 0.077 | 0.00311682 | cluster3 | DLK1 |
| 3.83E-13 | 1.47452943 | 0.58 | 0.247 | 1.27E-08 | cluster3 | HILPDA |
| 1.09E-24 | 1.41383837 | 1 | 0.972 | 3.61E-20 | cluster3 | FTL |
| 1.05E-20 | 1.38190074 | 0.71 | 0.267 | 3.47E-16 | cluster3 | NEAT1 |
| 1.28E-05 | 1.33699959 | 0.145 | 0.042 | 0.42376874 | cluster3 | TRH |
| 2.97E-35 | 1.32727873 | 0.377 | 0.049 | 9.83E-31 | cluster3 | GDF15 |
| 4.58E-12 | 1.27363895 | 0.739 | 0.432 | 1.51E-07 | cluster3 | HSPA5 |
| 9.98E-22 | 1.20421304 | 0.507 | 0.14 | 3.30E-17 | cluster3 | SLC2A1 |
| 1.39E-63 | 1.20040287 | 0.522 | 0.054 | 4.59E-59 | cluster3 | ADM |
| 1.46E-17 | 1.14892898 | 0.493 | 0.141 | 4.83E-13 | cluster3 | ANKRD37 |
| 4.71E-08 | 1.14282174 | 0.565 | 0.27 | 0.00155522 | cluster3 | TIMP1 |
| 2.08E-33 | 1.13115808 | 0.174 | 0.011 | 6.89E-29 | cluster3 | LINC02154 |
| 4.15E-29 | 1.10362465 | 0.681 | 0.188 | 1.37E-24 | cluster3 | SERPINH1 |
| 2.58E-14 | 1.0944657 | 0.507 | 0.173 | 8.51E-10 | cluster3 | DDIT3 |
| 4.59E-10 | 1.08793706 | 0.841 | 0.523 | 1.52E-05 | cluster3 | HSPB1 |
| 1.60E-16 | 1.07966968 | 0.797 | 0.429 | 5.28E-12 | cluster3 | P4HB |
| 0 | 2.849214 | 0.818 | 0.061 | 0 | cluster4 | TPH1 |
| 0 | 2.40849896 | 0.924 | 0.125 | 0 | cluster4 | GNB3 |
| 0 | 2.31759086 | 0.802 | 0.045 | 0 | cluster4 | GSG1 |
| 0 | 2.0111448 | 0.707 | 0.031 | 0 | cluster4 | PCAT4 |
| 0 | 1.88538292 | 0.831 | 0.117 | 0 | cluster4 | NEUROD1 |
| 0 | 1.88131969 | 0.594 | 0.114 | 0 | cluster4 | ISOC1 |
| 0 | 1.85498631 | 0.437 | 0.018 | 0 | cluster4 | RCVRN |
| 0 | 1.60505061 | 0.701 | 0.226 | 0 | cluster4 | PKIB |
| 3.80E-35 | 1.5256303 | 0.235 | 0.113 | 1.26E-30 | cluster4 | TCF7L2 |
| 0 | 1.4788523 | 1 | 0.601 | 0 | cluster4 | TTR |
| 0 | 1.45643274 | 0.41 | 0.032 | 0 | cluster4 | NRL |
| 0 | 1.416936 | 0.778 | 0.247 | 0 | cluster4 | PCBP4 |
| 0 | 1.41033522 | 0.583 | 0.045 | 0 | cluster4 | AANAT |
| 0 | 1.38231746 | 0.5 | 0.031 | 0 | cluster4 | AC004852.2 |
| 4.40E-58 | 1.36463645 | 0.216 | 0.068 | 1.46E-53 | cluster4 | CRABP1 |
| 2.60E-217 | 1.32479253 | 0.249 | 0.023 | 8.58E-213 | cluster4 | HOXB5 |
| 0 | 1.31796498 | 0.435 | 0.018 | 0 | cluster4 | ASMT |
| 0 | 1.95372251 | 0.838 | 0.057 | 0 | cluster5 | UBE2C |
| 6.81E-114 | 1.90182325 | 0.981 | 0.323 | 2.25E-109 | cluster5 | HMGB2 |
| 0 | 1.74434514 | 0.818 | 0.054 | 0 | cluster5 | TOP2A |
| 7.15E-292 | 1.73962175 | 0.838 | 0.064 | 2.36E-287 | cluster5 | NUSAP1 |
| 4.59E-37 | 1.71139277 | 0.714 | 0.362 | 1.52E-32 | cluster5 | HIST1H4C |
| 3.16E-122 | 1.66703573 | 0.773 | 0.139 | 1.04E-117 | cluster5 | PTTG1 |
| 3.42E-194 | 1.56147918 | 0.812 | 0.091 | 1.13E-189 | cluster5 | PCLAF |
| 0 | 1.53690472 | 0.831 | 0.052 | 0 | cluster5 | BIRC5 |
| 1.19E-200 | 1.42058093 | 0.682 | 0.06 | 3.94E-196 | cluster5 | CENPF |
| 1.15E-140 | 1.40704609 | 0.545 | 0.054 | 3.79E-136 | cluster5 | CCNB1 |
| 7.22E-180 | 1.37577822 | 0.721 | 0.077 | 2.39E-175 | cluster5 | CDK1 |
| 5.52E-77 | 1.37258482 | 1 | 0.815 | 1.82E-72 | cluster5 | HMGN2 |
| 2.04E-78 | 1.37153668 | 0.87 | 0.284 | 6.74E-74 | cluster5 | CKS2 |
| 4.50E-81 | 1.337139 | 0.994 | 0.828 | 1.49E-76 | cluster5 | H2AFZ |
| 1.88E-103 | 1.26719454 | 0.831 | 0.19 | 6.21E-99 | cluster5 | CKS1B |
| 1.14E-165 | 1.26544685 | 0.799 | 0.105 | 3.76E-161 | cluster5 | MAD2L1 |
| 1.47E-124 | 1.2143163 | 0.727 | 0.114 | 4.87E-120 | cluster5 | TYMS |
| 1.92E-78 | 1.21087008 | 1 | 0.918 | 6.34E-74 | cluster5 | TUBA1B |
| 0 | 1.18294758 | 0.74 | 0.038 | 0 | cluster5 | PBK |
| 8.03E-271 | 1.16490459 | 0.61 | 0.033 | 2.65E-266 | cluster5 | RRM2 |
| 3.92E-126 | 1.15889564 | 0.779 | 0.13 | 1.30E-121 | cluster5 | SMC4 |
| 1.22E-258 | 1.15852833 | 0.701 | 0.048 | 4.05E-254 | cluster5 | ZWINT |
| 2.85E-95 | 1.14469334 | 0.792 | 0.177 | 9.41E-91 | cluster5 | UBE2T |
| 2.78E-159 | 1.135325 | 0.766 | 0.102 | 9.18E-155 | cluster5 | CENPW |
| 4.62E-268 | 1.11092023 | 0.675 | 0.042 | 1.53E-263 | cluster5 | CDKN3 |
| 0 | 1.85095662 | 0.981 | 0.416 | 0 | cluster6 | CLU |
| 0 | 1.80687564 | 0.858 | 0.269 | 0 | cluster6 | APOE |
| 0 | 1.77451893 | 0.737 | 0.102 | 0 | cluster6 | S100B |
| 0 | 1.77241799 | 0.654 | 0.079 | 0 | cluster6 | SPARCL1 |
| 7.89E-133 | 1.77034541 | 0.9 | 0.567 | 2.61E-128 | cluster6 | PTN |
| 0 | 1.71245746 | 0.848 | 0.262 | 0 | cluster6 | SERPINF1 |
| 6.63E-200 | 1.65413004 | 0.826 | 0.302 | 2.19E-195 | cluster6 | IGFBP7 |
| 0 | 1.65145324 | 0.734 | 0.091 | 0 | cluster6 | TPPP3 |
| 0 | 1.65035788 | 0.529 | 0.036 | 0 | cluster6 | PMEL |
| 2.12E-121 | 1.56511455 | 0.394 | 0.098 | 7.01E-117 | cluster6 | FRZB |
| 0 | 1.50194507 | 0.775 | 0.193 | 0 | cluster6 | PTGDS |
| 1.48E-87 | 1.4996733 | 0.773 | 0.53 | 4.90E-83 | cluster6 | GPM6B |
| 0 | 1.43103974 | 0.789 | 0.145 | 0 | cluster6 | CD9 |
| 0 | 3.3411707 | 0.93 | 0.096 | 0 | cluster7 | COL3A1 |
| 0 | 3.29294471 | 0.946 | 0.064 | 0 | cluster7 | COL1A1 |
| 0 | 3.15538894 | 0.898 | 0.107 | 0 | cluster7 | MGP |
| 0 | 3.12872428 | 0.958 | 0.095 | 0 | cluster7 | COL1A2 |
| 0 | 2.83503457 | 0.866 | 0.06 | 0 | cluster7 | LUM |
| 8.78E-277 | 2.81975055 | 0.997 | 0.294 | 2.90E-272 | cluster7 | LGALS1 |
| 0 | 2.09977045 | 0.677 | 0.068 | 0 | cluster7 | SFRP2 |
| 8.38E-256 | 2.09386292 | 0.572 | 0.06 | 2.77E-251 | cluster7 | DLK1 |
| 5.33E-301 | 2.03943522 | 0.882 | 0.148 | 1.76E-296 | cluster7 | DCN |
| 5.64E-258 | 1.99926688 | 0.942 | 0.231 | 1.86E-253 | cluster7 | SPARC |
| 0 | 1.96272571 | 0.492 | 0.029 | 0 | cluster7 | FN1 |
| 3.02E-240 | 1.88863403 | 0.949 | 0.247 | 9.99E-236 | cluster7 | TIMP1 |
| 1.32E-225 | 1.83809911 | 0.962 | 0.28 | 4.36E-221 | cluster7 | IFITM3 |
| 0 | 1.77740973 | 0.751 | 0.028 | 0 | cluster7 | IGF2 |
| 0 | 1.77484746 | 0.895 | 0.092 | 0 | cluster7 | S100A11 |
| 0 | 1.63948492 | 0.78 | 0.069 | 0 | cluster7 | PCOLCE |
| 3.96E-187 | 1.60803356 | 0.859 | 0.244 | 1.31E-182 | cluster7 | FBLN1 |
| 4.04E-166 | 1.6059821 | 0.863 | 0.251 | 1.33E-161 | cluster7 | FOS |
| 0 | 1.59750701 | 0.626 | 0.012 | 0 | cluster7 | OGN |
| 0 | 1.50866457 | 0.581 | 0.031 | 0 | cluster7 | ITM2A |
| 0 | 1.50031561 | 0.754 | 0.077 | 0 | cluster7 | MFAP4 |
| 0 | 1.4542692 | 0.562 | 0.012 | 0 | cluster7 | POSTN |
| 0 | 1.4422772 | 0.735 | 0.02 | 0 | cluster7 | BGN |
| 8.34E-150 | 1.43640991 | 0.837 | 0.251 | 2.76E-145 | cluster7 | MEST |
| 0 | 1.42890322 | 0.665 | 0.06 | 0 | cluster7 | S100A10 |

**Table S3. List of cell stress-associated biological pathways**

| Term_ID | Description |
| --- | --- |
| GO:0034063 | stress granule assembly |
| GO:1903608 | protein localization to cytoplasmic stress granule |
| GO:0090400 | stress-induced premature senescence |
| GO:0043620 | regulation of DNA-templated transcription in response to stress |
| GO:0097201 | negative regulation of transcription from RNA polymerase II promoter in response to stress |
| GO:0033555 | multicellular organismal response to stress |
| GO:0043555 | regulation of translation in response to stress |
| GO:0036473 | cell death in response to oxidative stress |
| GO:0031098 | stress-activated protein kinase signaling cascade |
| GO:0006979 | response to oxidative stress |
| GO:0032055 | negative regulation of translation in response to stress |
| GO:0034976 | response to endoplasmic reticulum stress |
| GO:0035617 | stress granule disassembly |
| GO:0080134 | regulation of response to stress |
| GO:1902884 | positive regulation of response to oxidative stress |
| GO:0070059 | intrinsic apoptotic signaling pathway in response to endoplasmic reticulum stress |
| GO:1902882 | regulation of response to oxidative stress |
| GO:0070303 | negative regulation of stress-activated protein kinase signaling cascade |
| GO:0036475 | neuron death in response to oxidative stress |
| GO:0097165 | nuclear stress granule |

**Table S4. List of primary and secondary antibodies**

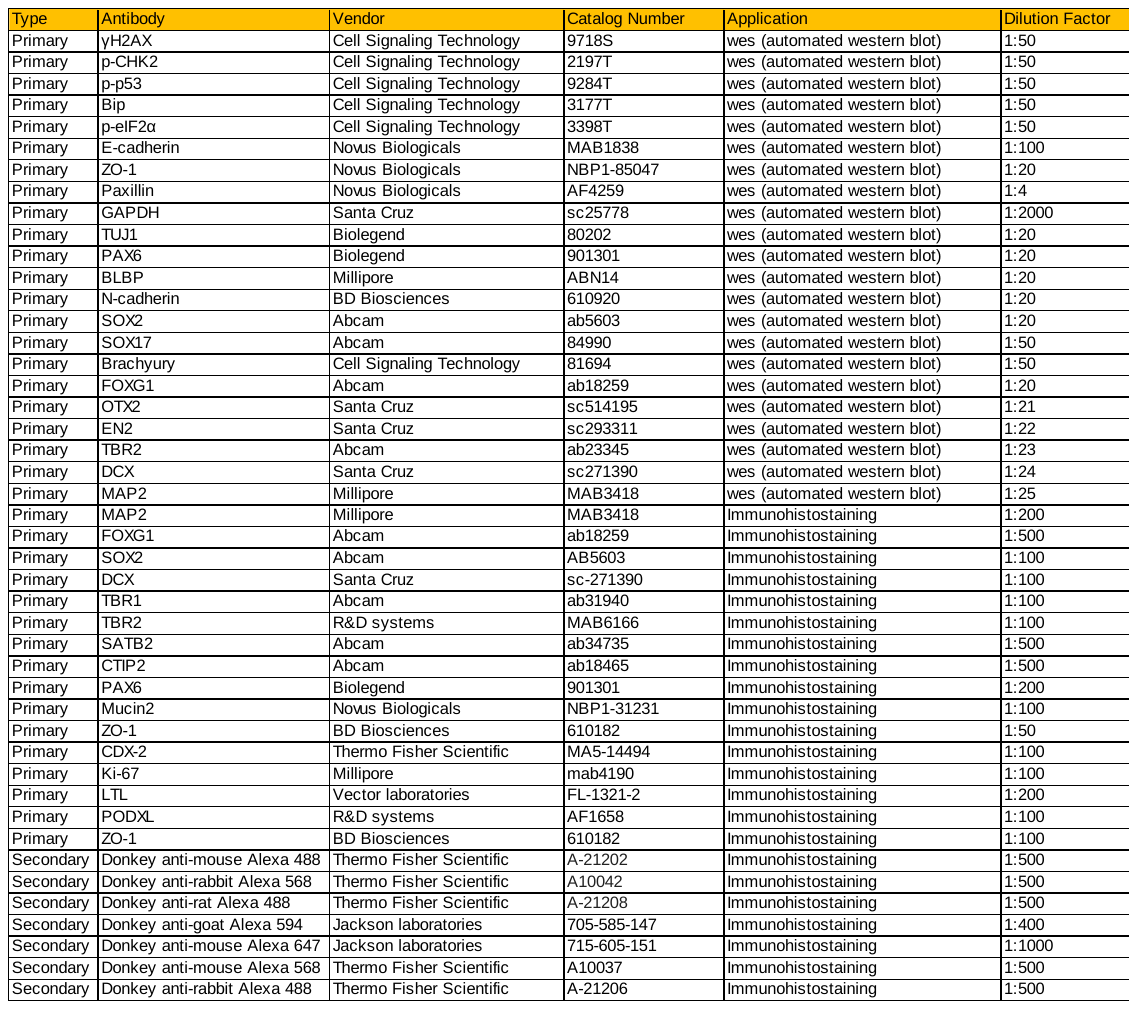

**SUPPLEMENTAL INFORMATION**

**Chemical compounds**

ROCK inhibitor Y-27632 (Tocris #1254) was used at 10 µM concentration, however, in some experiments at 20 µM to follow published protocols. To prepare the CEPT cocktail, we used the following concentrations: 50 nM chroman 1 (MedChem Express #HY-15392, stock solution 500 µM), 5 µM emricasan (Selleckchem #S7775, stock solution 50mM), polyamine supplement (Sigma Aldrich #P8483) diluted 1:1,000, according to the manufacturer’s recommendation, and 0.7 µM trans-ISRIB (Tocris #5284, stock solution 7 mM). DMSO was used as the solvent for all the small molecule compounds following the manufacturer’s recommendation. More details on how to prepare the CEPT cocktail can be found here:

<https://ipsc.ncats.nih.gov/wp-content/uploads/2021/11/SCTL-CEPT-Protocol.pdf>

**Cell culture and brain organoid formation**

Human ESC (WA09) and iPSC lines (LiPSC-GR1.1 from NIH Common Fund; GM23279 and GM25256 were purchased from Coriell Institute for Medical Research) were maintained under feeder-free condition using mTeSR Medium (STEMCELL Technologies) and vitronectin (VN)-coated plates (Thermo Fisher Scientific). Cell lines were confirmed to be karyotypically normal and mycoplasma-free (MycoAlert Detection Kit, Lonza). Cells were routinely passaged when cultures reached 70-90% confluency (every 3 to 4 days) using 0.5 mM EDTA diluted in phosphate buffered saline (PBS) without calcium or magnesium (Thermo Fisher Scientific). Cerebral organoids were generated using a commercial kit (STEMdiff™ Cerebral Organoid Kit, STEMCELL Technologies). In brief, low-passage hPSCs were detached using Accutase (Thermo Fisher Scientific) and plated in 96-well ultra-low attachment (ULA) round-bottom plates (Corning) at a density of 9,000 cells per well in 100 µl of EB Formation Medium (STEMdiff™ Cerebral Organoid Kit) to generate EBs with either the addition of 10 µM Y-27632 or the CEPT cocktail. An additional 100 µL of EB Medium was added every other day for 5 days. At day 4-6, or typically when EBs showed round, smooth edges and reached between 400-600 µm in diameter, each EB was transferred into each well of 24-well ultra-low attachment plate (Corning) in 0.5 mL of Neural Induction Medium (STEMdiff™ Cerebral Organoid Kit). At day 6-8, when smooth translucent edges of the neuroepithelium emerged, each organoid was embedded in 15 µL droplet of Matrigel (Corning). Droplets of Matrigel with the organoids were then transferred into a 6-well ULA plate (Corning) in Neural Expansion Medium (STEMdiff™ Cerebral Organoid Kit). After 3 days, the medium was changed to Neural Maturation Medium (STEMdiff™ Cerebral Organoid Kit). The plate of organoids was placed on an orbital shaker. The medium was changed every 3 days. To generate cerebral organoids based on a previously developed protocol by Velasco et al., WA09 cell lines were dissociated into single cells using Accutase (Thermo Fisher Scientific) and plated in 96-well ULA round-bottom plates (Corning) at a density of 6,000 cells per well with either the addition of 20 µM Y-27632 or the CEPT cocktail in cortical differentiation medium (CDM) I, containing Glasglow-MEM (Gibco), 20% Knockout Serum Replacement (Gibco), 0.1 mM Minimum Essential Medium non-essential amino acids (Gibco), 1 mM Sodium Pyruvate (Gibco), and 0.1 mM 2-mercaptoethanol (Gibco). TGFβ inhibitor SB431542 (Tocris) and WNT inhibitor IWR1 (Tocris) were added at a concentration of 5 µM and 3 µM, respectively. Media was changed every 3 days with CDM1 48 hours after seeding. After 18 days, the aggregates were cultured in 100-mm ULA culture dishes (Corning) on an orbital shaker and media was changed to CDM II, containing DMEM/F12 medium (Gibco), 2 mM Glutamax (Gibco), 1% N2 (Gibco), and 1% Chemically Defined Lipid Concentrate (Gibco). Media was changed every 3 days until day 35. All cultures were maintained at 37 °C under humidified 5% CO_2_ and atmospheric O_2_.

**EB viability analysis**

Phase-contrast images of EBs were obtained with an Incucyte Zoom Live Cell Analysis System (Sartorius). The CellTiter-Glo 3D assay (Promega G9681) was carried out following the manufacturer’s instructions to quantify cell viability. Luminescence signal was read using the PHERAstar FXS microplate reader (BMG LABTECH). EB size was measured using the Celigo Imaging Cytometer (Nexcelom Biosciences). Live and dead cells were stained using a two-color fluorescence live/dead assay kit (Thermo Fisher Scientific, 1:2,000) and imaged with an DMi8 microscope (Leica) or an Opera Phenix high-content microscope (PerkinElmer) using appropriate filters. During each assessment, single EBs were cultured in 96-well ULA round-bottom plates (Corning).

**Western blot**

The Wes automated western blotting system (ProteinSimple) was used following the manufacturer’s instructions. Briefly, cell lysate and reagents were loaded into assay plates and placed into the Wes system, which automatically loads cell lysates into the capillary for electrophoretic protein separation. Proteins of interest were identified using primary antibodies, HRP-conjugated secondary antibodies followed, and chemiluminescent substrate. Chemiluminescent signals were detected by a camera integrated into the Wes system and quantified by using the Compass software. All western blot data are displayed by lanes in virtual blot-like images. Detailed information on primary and secondary antibodies is provided in Supplementary Table S4.

**Individual EB culture in E6 medium and RASL-Seq**

For EB formation, WA09 hESCs were single-cell dissociated using Accutase (Thermo Fisher Scientific) and plated into 96-well ULA round-bottom plates (Corning) at a density of 20,000 cells per well in Essential 6 (E6) Medium (Thermo Fisher Scientific) for 7 days. Three different protocols were used to differentiate hESCs into ectoderm, mesoderm, and endoderm, which served as positive controls for lineage-specific gene expression. For neural induction, hESCs were treated with LDN-193189 (100 nM; Sigma Aldrich) and A83-01 (2 µM; Tocris) in E6 Medium for 6 days. For either mesoderm or endoderm induction, commercially available kits were used for 5 days following the instructions of the manufacturer (STEMdiff™ Mesoderm Induction Medium and STEMdiff™ Definitive Endoderm Kit, STEMCELL Technologies). Samples were harvested and lysed in TCL lysis buffer (Qiagen).

RASL-Seq was carried out using an automated liquid handler (Biomek i7, Beckman Coulter) equipped with a multichannel head and a temperature-controlled BioRad C1000 thermal cycler (Biorad). Lysates were transferred to TurboCapture 384 format plates (Qiagen) and incubated at RT 1 hour at 100 rpm for polyA RNA capture. The cell lysate was then washed with a binding buffer (20 mM Tris-HCl, pH 7.5 at 25⁰C, 500 mM NaCl, 1 mM EDTA, 0.1% SDS) twice and 20 µL of probes mixture was added into each well with 0.05 pmol/oligo in binding buffer. The binding mixture was incubated at 65⁰C for 10 minutes and gradually the temperature was lowered to 45⁰C by at a 3 degree/per minute rate before 1 hour incubation at 45 ⁰C in the thermocycler. The ligation step was performed after three washes with wash buffer (20 mM Tris-HCl, pH7.5 at 25 ⁰C, 100 mM NaCl, 1 mM EDTA, 0.1% Tween 80) and one wash with ligation buffer (40 mM Tris-HCl, pH7.5 at 25 ⁰C, 10 mM MgCl2, 10 mM DTT, 10 µM ATP). A 20 µL ligation mixture containing 5U (Weiss unit) of T4 DNA ligase in ligation buffer was added into each well. After ligation at 37⁰C for 1 hour, the wells were washed three times with wash buffer. The final products were eluted with 25 µl elution buffer. Final products of 5 µL/sample were used for PCR amplification with 0.05 µM barcoded primers and AmpliTaqGold DNA polymerase (0.06 U/reaction, Thermo Fisher Scientific) in buffer I supplied by the manufacturer. The dual indexed PCR products were then pooled as one single sample and purified using size-selected column to remove primer dimers (Zymo Research). The purified samples were quantified and sequenced on an Illumina NextSeq 550 using custom sequencing primers.

RASL-Seq data processing and counting were performed using the computational resources of the NIH HPC Biowulf cluster (http://hpc.nih.gov). RASL-Seq libraries were 8 bases-dual indexed at the PCR amplification step. Each well has a unique pair of sequence barcodes to identify the corresponding treatment.

RASL-Seq raw data demultiplexing was accomplished using Illumina BaseSpace automation. Genome alignment and read counting used a custom subset of the hg38 reference genome since custom oligo-dt primers were used. A custom python pipeline was used for alignment based on Hamming distance. Details on the method can be found at https://github.com/ncats/raslseq. Downstream analysis of RASL-Seq reads was done similarly as with bulk RNA-Seq, using DESeq2. Probes of which the maximum responding read count were less than 20 (maxBM: average normalized count between treatment and basal condition) across all conditions were removed from analysis to minimize the noise.

**High-content imaging and analysis of organoids**

Cerebral organoids (hESC WA09) were generated at varying cell numbers (1,500-7,500 cells per well) and fixed with 4% PFA at day 10, which represents the end of neuroepithelial expansion phase according to the kit-based protocol (STEMCELL Technologies). Organoids were washed with DPBS and nuclei were stained with Hoechst 33342 solution (Thermo Fisher Scientific, 1:2000 in DPBS) for 30 min at room temperature. Organoids were washed three times with DPBS and individual organoids were placed into a 96-well ULA round-bottom plate (Corning) immersed in ScaleS4 clearing solution, which is composed of 40 w/v% D-(-)-sorbitol (Sigma), 10 w/v% glycerol (Sigma), 4 M Urea (Sigma), 0.2 w/v% Triton X-100 (Sigma), and 15 v/v% DMSO (Sigma) in ultrapure water (Gibco). The plate containing organoids were incubated in clearing solution for 24 hours at 37°C on a shaker. Following clearing, the organoids were imaged on the Opera Phenix high-content imaging microscope (PerkinElmer) at excitation/emission = 470-495/515-575. A customized image analysis script for automated measurement of neurosphere features was developed and executed in the Matlab scientific computing environment (Matlab version 2020b, The Mathworks, Inc.). Briefly, algorithmic local thresholds and binary morphological region operations segment the total spheroid volume and neurosphere center volumes from the XYZ digital image stack. The total neurosphere volume external dimensions define the hemisphere size of the structure. Hemisphere volume is used to approximate full size volume and allows measurement of hemisphere features even when the entire neurosphere cannot be analyzed because the total z-size is too large for acquisition. Neurosphere volume without neurosphere centers is calculated for the automatically defined hemisphere volume. Detection of individual nuclei is performed by local maximal intensity in 3D space.

**Bulk RNA sequencing libraries preparation, sequencing reaction, and analysis**

RNA was extracted (three samples for each group, each consisting of three organoids) using the RNeasy Mini Kit (Qiagen). For reproducibility studies, RNA was extracted from single organoids. RNA was quantified using the Agilent RNA 600 Kit (Agilent) on a BioAnalyzer 2100 (Agilent). RNA (500ng, RIN >8 was used to prepare RNA-seq libraries with the TruSeq Stranded mRNA Library Prep Kit (Illumina) according to the manufacturer’s protocol. Sequencing libraries were quantified by PCR using the KAPA library quantification kit for Illumina platforms (KAPA Biosystems) using QuantStudio 12K Flex Real-Time PCR System (Thermo Fisher Scientific). All samples were normalized according to concentration and pooled. Libraries were loaded and sequenced using the Illumina NextSeq 550 system. Bioinformatics analysis for bulk RNA-seq was carried out using the computational resources of the NIH HPC Biowulf cluster (http://hpc.nih.gov) using R language 3.6.0 (https://cran.r-project.org/). Bulk RNA-Seq samples were quality trimmed using Trimmomatic 0.36 and the TruSeq3 paired-end adapters. STAR aligner 2.7.6a followed by HTSeq-count 0.9.1 produced deduplicated counts of reads in genes. UCSC gene counts were normalized using the default median-of-ratios method in DESeq2 1.24.0. Differential expression tests used the lfcShrink function and gene set enrichment was performed using Enrichr API package enrichR 1.0. Gene set libraries queried were the Gene Ontology Biological Process 2018 and ARCHS4 Tissues Database.

**Correlation analysis of individual organoids**

For the generation of correlation plots, a correlation matrix of Pearson R values across all transcripts between all pairs of samples shown in each plot was generated using the cor base R function based on raw counts. R2 values were computed from the correlation matrix and passed to the corrplot function from the corrplot R package (version 0.84) to create the correlation plot. For the silhouette plot, a distance matrix was generated by subtracting Pearson R values (computed in the same way as the correlation plot prior to squaring) from 1. Samples were split into groups based on cell line and treatment as indicated by the gray and blue shaded rectangles. For each group, a medoid sample was identified based on the highest average Pearson correlation with other members of that group. The distance matrix and medoid identities were passed to the sil function from the kmed R package (version 0.3.0) to create the silhouette plot.

**Histological analysis**

Organoids were fixed in 4% PFA (Thermo Fisher Scientific) at room temperature for 20 min, washed with PBS three times, and incubated in 30% sucrose at 4°C overnight. Tissues were embedded in O.C.T. compound (Fisher Scientific), cut into 20-µm sections, and mounted on microscope slides (Fisher Scientific) for staining. For histological analysis, sections were stained with H&E, dehydrated in ethanol and xylene, and coverslipped using Permount mounting medium (Fisher Scientific). For immunohistochemical analysis, sections were permeabilized and blocked with 0.3% Triton X-100 and 5% BSA in PBS for 1h. Detailed information on primary and secondary antibodies is provided in Supplementary Table S4. Slides were mounted with ProLong Glass Antifade Mountant with NucBlue Stain (Thermo Fisher Scientific). Fluorescence images were taken with the Zeiss LSM 710 confocal microscope using appropriate filters.

**Single-cell RNA sequencing libraries preparation, sequencing reaction, and analysis**

Cerebral organoids (hESCs WA09) were cultured until day 72, washed twice in DPBS (Thermo Fisher Scientific), and dissociated into a single-cell suspension using Embryoid Body Dissociation Kit (Miltenyi Biotec) and gentleMACS Dissociator (Miltenyi Biotech) in gentleMACS C Tube (Miltenyi Biotec) following the manufacturer’s protocol. The cell suspension was filtered through a 70-μm strainer (Miltenyi Biotec) to remove cell debris and clumps. The strained cell suspension was centrifuged at 300 g for 5 min and resuspended in DPBS to obtain approximately 5,000 cells/sample/lane of a 10x microfluidic chip device (10X Genomics). This suspension was kept on ice for no longer than 30 min. The microfluidic chip device was loaded onto a Chromium™ Single Cell 3’ Chip (10X Genomics) and processed through the Chromium Controller to generate single-cell GEMs (Gel Beads in Emulsion). Libraries were prepared from the single-cell GEMs with the Chromium™ Single Cell 3’ Library & Gel Bead Kit v2 (10x Genomics) according to the manufacturer’s protocol. Libraries from different samples were pooled together based on molar concentrations and sequenced on a NextSeq 550 instrument (Illumina) with 26 bases for read 1, 98 bases for read 2, and 8 bases for Index 1.

The Cellranger software package from 10X Genomics, Inc. (version 3.0.1) was used to process raw BCL files from single-cell sequencing as follows. Pipeline details can be found at https://github.com/cemalley/Ryu_cerebral_organoids. Demultiplexing and FASTQ generation were done with the mkfastq command, and the count command created gene expression matrices. Dense matrices were created with the mat2csv command. Embryonic stem cell and iPSC lines were analyzed in the Seurat R package (Seurat 2.3.4; R 3.5.2) (Stuart et al., 2019). Unbiased clustering within samples was performed using FindClusters, which were then refined based on expression differences between clusters (Figure 6A). Data visualizations were made in R and with the ggplot2 package (3.1.0). Gene expression enrichment analysis was performed with the same methods as for bulk RNA-Seq.

**Intestinal and kidney organoids formation**

Intestinal organoids were generated using a commercial kit (STEMdiff™ Intestinal Organoid Kit, STEMCELL Technologies). Briefly, on day -2, either hESCs (WA09) or hiPSCs (LiPSC-GR1.1) previously maintained in mTeSR1 (STEMCELL Technologies) were detached using 0.5 mM EDTA in PBS without calcium or magnesium (Thermo Fisher Scientific). The detached cell clumps (50-200 µm in diameter) were plated in Matrigel (Corning)-coated 24-well plate (Corning) at densities of approximately 6,000 clumps per well in 0.5 mL mTeSR1 with either the addition of 10 µM Y-27632 or CEPT cocktail for the first 24 hours after cell seeding. On day 0, media was switched to 0.5 mL/well of Definitive Endoderm Differentiation Medium (STEMdiff™ Intestinal organoid Kit) with daily media change. On days 3-9, media was switched to 0.5 mL/well of Mid-/Hindgut Differentiation Medium (STEMdiff™ Intestinal organoid Kit) with daily media change. Released spheroids from the monolayer culture were collected and embedded in Matrigel for further maturation into small intestinal organoids. Free-floating spheroids (~50) were embedded in 50 µl cold Matrigel and transferred to a 24-well Nunclon Delta surface plate (Thermo Fisher Scientific). After 20 min incubation at 37°C to allow the Matrigel to solidify, 0.5 mL of pre-warmed Intestinal Organoid Growth Medium (STEMdiff™ Intestinal organoid Kit) was added. Media was changed every 3-4 days. The organoids were passaged every 7-10 days. On day 28, cells were fixed and processed for histological analysis (see above).

Kidney organoids were generated using a commercial kit (STEMdiff™ Kidney Organoid Kit, STEMCELL Technologies). Briefly, on day -3, either hESCs (WA09) or hiPSCs (LiPSC-GR1.1) previously maintained in mTeSR1 (STEMCELL Technologies) were dissociated using Accutase (Thermo Fisher Scientific) and plated into Matrigel (Corning)-coated 96-well µ-plates (Ibidi) at densities of 500 cells per well in 200 µL of mTeSR1 with either the addition of 10 µM Y-27632 or CEPT cocktail. On day -2, medium was removed and cells were overlayed with 100 µL of cold mTeSR1 supplemented with Matrigel (0.25 mg/mL). On day -1, media was removed and changed with 100 µL of mTeSR1. On day 0, medium was switched to 100 µL of Stage 1 Kidney Organoid Differentiation Medium (STEMdiff™ Kidney organoid Kit). On day 1.5, medium was switched to 100 µL of Stage 2 Kidney Organoid Differentiation Medium (STEMdiff™ Kidney organoid Kit). From days 4 to 18, Stage 2 Medium was replaced every 2–3 days. On day 18, cells were fixed with 4% PFA for 15 min, washed with PBS for 15 min, and blocked with 10% donkey serum (Jackson Laboratories) in PBS for 1 hour. Detailed information on primary and secondary antibodies is provided in Supplementary Table S4. Nuclei were stained with Hoechst 33342 (Thermo Fisher Scientific, 1:2000). Fluorescence images were taken with the Zeiss LSM 710 confocal microscope using appropriate filters.

**Statistical analysis**

All results are shown as mean ± S.D. Statistical tests included unpaired, two-tailed Student’s t-tests and one-way ANOVA for multiple comparisons with Tukey’s significant difference post hoc test using GraphPad Prism 9.0.0. A *P* value of <0.05 was considered statistically significant. *P* values denote as **p*<0.05, ***p*<0.01, ****p*<0.001, or *****p*<0.0001.
